# Transfer function for YAP/TAZ nuclear translocation revealed through spatial systems modeling

**DOI:** 10.1101/2020.10.14.340349

**Authors:** Kiersten E. Scott, Stephanie I. Fraley, Padmini Rangamani

## Abstract

YAP/TAZ is a master regulator of mechanotransduction whose functions rely on translocation from the cytoplasm to the nucleus in response to diverse physical cues. Substrate stiffness, substrate dimensionality, and cell shape are all input signals for YAP/TAZ, and through this pathway, regulate critical cellular functions and tissue homeostasis. Yet, the relative contributions of each biophysical signal and the mechanisms by which they synergistically regulate YAP/TAZ in realistic tissue microenvironments that provide multiplexed input signals remains unclear. For example, in simple 2D culture, YAP/TAZ nuclear localization correlates strongly with substrate stiffness, while in 3D environments, YAP/TAZ translocation can increase with stiffness, decrease with stiffness, or remain unchanged. Here, we develop a spatial model of YAP/TAZ translocation to enable quantitative analysis of the relationships between substrate stiffness, substrate dimensionality, and cell shape. Our model couples cytosolic stiffness to nuclear mechanics to replicate existing experimental trends, and extends beyond current data to predict that increasing substrate activation area through changes in culture dimensionality, while conserving cell volume, forces distinct shape changes that result in nonlinear effect on YAP/TAZ nuclear localization. Moreover, differences in substrate activation area versus total membrane area can account for counterintuitive trends in YAP/TAZ nuclear localization in 3D culture. Based on this multiscale investigation of the different system features of YAP/TAZ nuclear translocation, we predict that how a cell reads its environment is a complex information transfer function of multiple mechanical and biochemical factors. These predictions reveal design principles of cellular and tissue engineering for YAP/TAZ mechanotransduction.

**STATEMENT OF SIGNIFICANCE:** In chemical engineering, a transfer function is a mathematical function that models the output of a reactor for all possible inputs, and enables the reliable design and operation of complex reaction systems. Here, we apply this principle to cells to derive the transfer function by which substrate stiffness is converted into YAP/TAZ nuclear localization. This function is defined by a spatial model of the YAP/TAZ mechano-chemical sensing network, wherein key spatial and physical inputs to the system, namely cell and nuclear shape, surface area to volume ratios of cytoplasmic and nuclear compartments, substrate dimensionality, substrate activation area, and substrate stiffness, are all integrated. The resulting model accounts for seemingly contradictory experimental trends and lends new insight into controlling YAP/TAZ signalling.

## INTRODUCTION

The interplay between cell shape, cell-substrate interaction, and nuclear translocation of transcription factors is critical for cellular homeostasis; misregulation of these relationships can often result in pathological states. An important example of this is the regulation of nuclear translocation of YES-associated protein 1 (YAP1) and TAZ (YAP/TAZ), one of the primary sensors of the cell’s mechanical states. YAP/TAZ is a co-transcriptional activator repressed in the Hippo tumor suppressor pathway, which regulates proliferation and cell differentiation to maintain organ size (1). While YAP/TAZ has been implicated as a key regulator of mechanotransduction by translating extracellular matrix cues, specifically matrix stiffness, it remains unclear how other biophysical and biochemical features of the cell interact with substrate stiffness to initiate downstream signaling events all the way to nuclear translocation (2, 3) (Fig. 1A).

**Figure 1.**
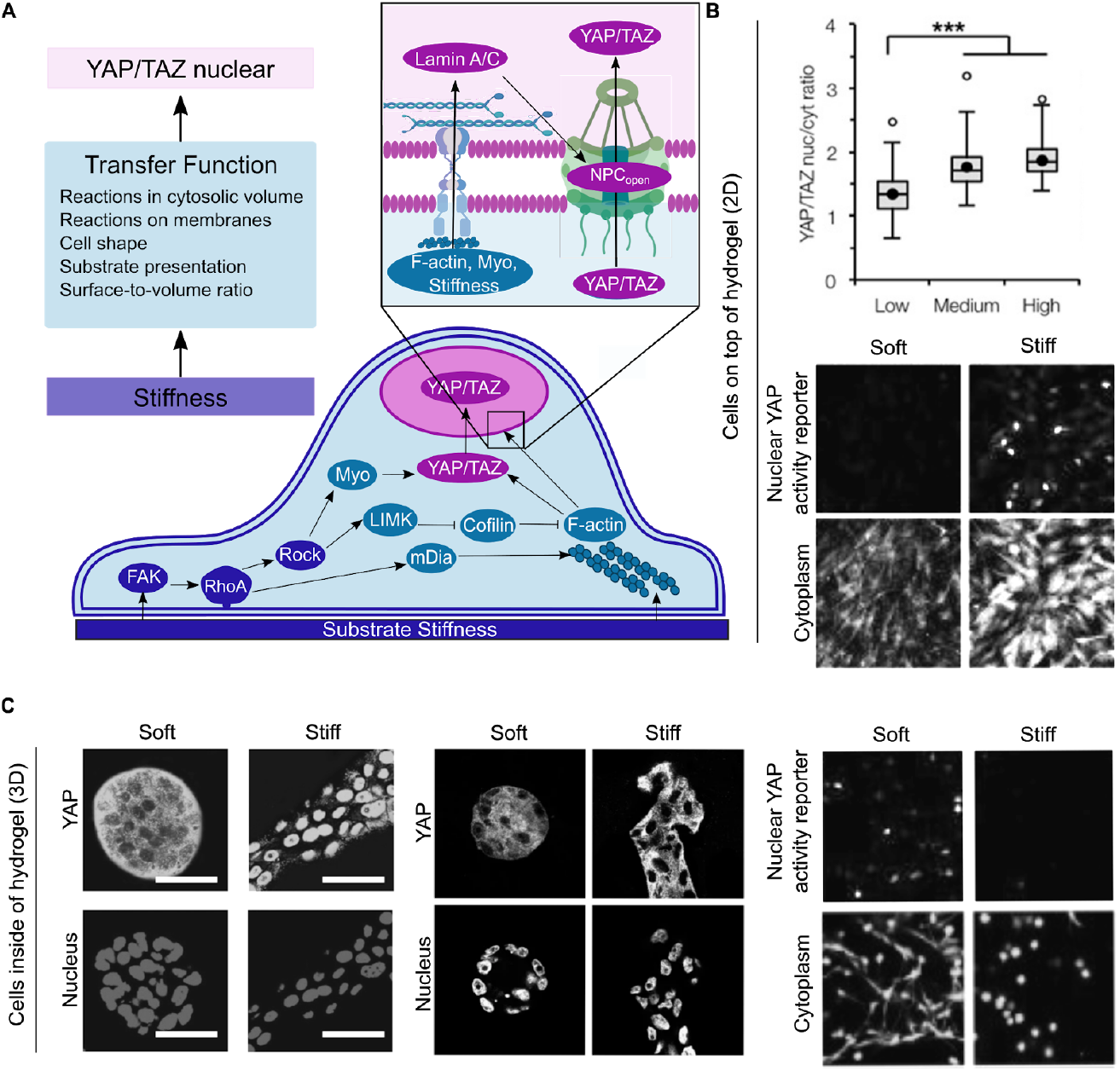
(*A*) The transfer function by which substrate stiffness is converted into YAP/TAZ nuclear localization is defined by a spatial model of the YAP/TAZ mechano-chemical sensing network. Reactions that respond to substrate stiffness are carried out near the substrate-adjacent plasma membrane of the cell by integrins and propagated by FAK, RhoA, and ROCK (components in blue, Module 1 of model). These signals are translated into cytoskeleton regulation by mDia, LIMK, Cofilin, Myosin, and F-actin (components in teal, Module 2 of model). The cytoskeletal regulators then impinge on YAP/TAZ phosphorylation state, regulating the ability of YAP/TAZ to enter the nucleus (components in pink, Module 3 of model). (Inset) The actin cytoskeleton also directly regulates YAP/TAZ nuclear import by transferring cytoskeletal stiffness to the nucleus by way of Lamin A/C, stretching nuclear pore complexes (17). (*B*) Cells cultured in 2D conditions, on top of substrates, respond to increasing stiffness by increasing YAP localization to the nucleus. Data adapted with permission from (8) and (29). (*C*) Cells cultured in 3D conditions, inside of substrates, respond to increasing stiffness by increasing YAP nuclear localization (left), not changing YAP localization (middle), or decreasing YAP nuclear localization (right). Data adapted with permission from(9), (30), and (29).

Extracellular matrix stiffness is a key environmental input, instructing the cell about its pericellular space (4). However, the cell must interpret this input by regulating integrin clustering (5) and through cytoskeletal contractility including both myosin and actin activities (6) through actin binding proteins (7). Such interpretation likely involves sensing the dimensionality of stiffness cues through the spatial organization of the extracellular substrate as well as spatial restrictions on cell spreading imposed by the substrate format. Specifically the presentation of two-dimensional (2D) versus three-dimensional (3D) extracellular matrix structure has revealed a critical need to unravel the information transfer function underlying the context-dependence of YAP/TAZ nuclear translocation (Fig. 1B,C). In 2D cell culture, nuclear YAP/TAZ localization increases with increasing substrate stiffness (Fig. 1B) (3, 8). However, in 3D, the trends of YAP/TAZ and stiffness are more complex (Fig. 1C). For example, while YAP/TAZ nuclear localization increased with increasing stiffness in 3D Matrigel/collagen matrices (9), no statistically significant relationship between YAP/TAZ nuclear translocation and stiffness was observed with physically crosslinked alginate hydrogels (10). Caliari *et al*. observed that in 3D tunable hyaluronic acid gels that YAP/TAZ nuclear localization does not correlate with stiffness but does depend on the degradability of the scaffold in 3D (8).

While these results seem confounding, it is important to note that several features of the substrate and cell change in tandem with substrate dimensionality. Key factors that can be altered between 2D and 3D cell culture systems include cell shape, cell-substrate interaction, and nuclear shape. Specifically, cell volume and shape change as cells spread on 2D substrates (11), but 3D substrates are inherently confining and can restrict volume and shape changes due to fewer physical degrees of freedom (12). By the same token, cell-substrate contact area changes between 2D, basal cell-ECM interactions, and 3D, basal and apical cell-ECM interactions, culture systems. When cells are cultured on 2D surfaces, cells display large focal adhesions (13, 14). Yet, cells grown in 3D soft matrix possess smaller, more nascent focal adhesions that diffuse not only in the basal side of the cell but across the apical surface of the cell (14). Additionally, nuclear shape and mechanical properties are also different between 2D and 3D. For example, in 2D, stress fibers link focal adhesions in the lamellipodia to LINC complexes at the cell nucleus (15). LINC (linker of nucleoskeleton and cytoskeleton) complexes are transmembrane protein complexes that bind lamin A in the inner nuclear membrane side to stress fibers on the cytoplasmic side; interaction of stress fibers with LINC complexes form a nuclear actin cap. In 3D, the number of stress fibers over the nucleus of the cell and the amount of lamin A are both lower (16). It has been suggested that the interaction of stress fibers and the LINC complex may facilitate nuclear flattening, which stretches nuclear pores to increase YAP/TAZ nuclear localization (17). Thus, YAP/TAZ nuclear localization is linked to mechanical properties of the matrix through multiple layers of spatiotemporal integration of mechanochemical information (Fig. 1A). Therefore, how information transfer through multiple mechanical, spatial, and temporal scales ultimately orchestrates YAP/TAZ nuclear translocation remains poorly understood.

Given the complexity of the inputs in such a mechanotransduction system, computational modeling offers a systematic way to conduct multivariate experiments and generate experimentally testable predictions. Computational modeling of signal transduction has played a key role in identifying the emergent properties in signaling networks (18–20). Subsequently, spatial modeling of cell shape and signaling has revealed the importance of geometry, surface-volume effects, and the localization of kinases in modulating signal transduction (21–25). More recently, substantial advances in modeling the interaction between mechanical cues and biochemical signaling have resulted in shedding light on this intricate coupling, paving the way for modeling efforts that can integrate different inputs to the cell (26, 27).

Specifically, modeling YAP/TAZ translocation by Sun and colleagues has revealed the dependence of YAP/TAZ response on stiffness as a function of multiple signaling pathways and the different molecule concentrations using a well-mixed model (28). Building on these technical and scientific advances, in this study we sought to answer the following questions: How does cell shape and culture dimensionality affect the nuclear translocation of YAP/TAZ for different substrate stiffnesses? Does substrate contact area play a role in modulating these relationships? And finally, how does nuclear shape affect YAP/TAZ nuclear localization under these different conditions? To answer these questions, we used a systems biophysics approach to computationally model the different scenarios that may affect YAP/TAZ mechanotransduction. By doing so, we sought to build a transfer function -- essentially, an input-output relationship, which can capture the impact of multiple cues on YAP/TAZ nuclear translocation. Indeed, simulations from the model predict that YAP/TAZ nuclear translocation is a complex information transfer function of cell shape, substrate contact area, plasma membrane area, and nuclear shape.

## METHODS

### Modular construction of the model

We developed a 3D spatial model of YAP/TAZ nuclear localization as a function of substrate stiffness (Fig. 1A). The modules are divided as follows: (1) substrate stiffness sensing by a biochemical cascade, (2) biophysical regulation of the cytoskeleton in response to substrate stiffness, and (3) YAP/TAZ nuclear localization as a function of cytoskeletal and nuclear properties. Each module in detail described below and the reactions and parameters are explained in detail in Tables S1, S2.

### Module 1: Substrate stiffness sensing

In this module (Fig. 1A, blue components), we describe the events that connect the stiffness of the extracellular matrix to the upstream signaling events, including the activation of focal adhesion kinase (FAK), RhoA activation, and Rho kinase (ROCK) activation. All of these events are adapted from the well-mixed model in Sun *et al. (28)* and converted to a spatial model.

#### 1. FAK activation

Stiffness of the extracellular matrix activates clustering of integrins and associated proteins which facilitates focal adhesion kinase phosphorylation (31). In this model, matrix stiffness is incorporated into the signaling pathway as a stimulus such that FAK is activated by the stiffness, *E* (Table 1, Rates 1 and 2), using kinetics similar to that of Michaelis-Menten kinetics (28, 32). The activation of FAK was implemented as a boundary condition at the plasma membrane in the spatial model using a position Boolean operator (see supplementary material for details).

**Table 1:** Partial differential equations for the events outlined in Module 1 along with species, locations, spatial equations and reaction rates used in the model. See Table S2 for the definition of the parameters.

| # | Species | Comp. | PDE<br>Boundary Condition | Reaction Rate |
| --- | --- | --- | --- | --- |
| 1 | $[FAK]$ | Cyto | $\frac{d[FAK]}{dt} = D_{[FAK]}\nabla^2[FAK] - R_2$ $-D_{FAK}(n \cdot \nabla[FAK])\vert _{PM} = -n_r R_1$ | $R_1 = k_f[FAK] + k_{sf}\frac{E}{C + E}[FAK]$ |
| 2 | $[FAK_p]$ | Cyto | $\frac{d[FAK_p]}{dt} = D\nabla^2[FAK_p] + R_2$ $-D_{FAK_p}(n \cdot \nabla[FAK_p])\vert _{PM} = n_r R_1$ | $R_2 = -k_{df}[FAK_p]$ |
| 3 | $[RhoA_{GDP}]$ | Cyto | $\frac{d[RhoA_{GDP}]}{dt} = D_{RhoA_{GDP}} \nabla^2 [RhoA_{GDP}]$ $-D_{RhoA_{GDP}}(n \cdot \nabla [RhoA_{GDP}])\vert _{PM} = -n_r R_3 - R_4$ | $R_3 = k_{fkp}(\gamma[FAK_P]^n + 1)[RhoA_{GDP}]$ |
| 4 | $[RhoA_{GTP}]$ | PM | $\frac{d[RhoA_{GTP}]}{dt} = D_{RhoA_{GTP}} \nabla^2 [RhoA_{GTP}] + n_r R_3 + R_4$ | $R_4 = -k_{d\rho}[RhoA_{GTP}]$ |
| 5 | $[ROCK]$ | Cyto | $\frac{d[ROCK]}{dt} = D_{ROCK} \nabla^2 [ROCK] - R_5$ $-D_{ROCK}(n \cdot \nabla [ROCK])\vert _{PM} = -R_6$ | $R_5 = -k_{drock}[ROCK_A]$ |
| 6 | $[ROCK_A]$ | Cyto | $\frac{d[ROCK_A]}{dt} = D_{[ROCK_A]} \nabla^2 [ROCK_A] + R_5$ $-D_{ROCK_A}(n \cdot \nabla [ROCK_A])\vert _{PM} = R_6$ | $R_6 = k_{r\rho}[RhoA_{GTP}][ROCK]$ |

#### 2. RhoA activation downstream of FAK

RhoA is known to be activated downstream of FAK (33) and is also upregulated with increasing rigidity of ECM in 3D environments (34). RhoA-GTP is modeled as a membrane species while RhoA-GDP is modeled as a cytosolic species, consistent with experimental observation (35). The reaction flux of RhoA is modeled as a function of phospho-FAK as shown in (Table 1, Rate 3) (28).

#### 3. RhoA activates Rho Kinase (ROCK)

ROCK is a downstream target of RhoA (36) and increases myosin activity by phosphorylating myosin light chain and inhibiting myosin phosphatase (37) (Table 1, Rates 4 and 5). The activation of ROCK by RhoA-GTP is modeled as a boundary flux at the plasma membrane. Inactivation of ROCK is modeled as a first order reaction in the cytoplasm.

### Module 2: Cytoskeleton regulation

In this module (Fig. 1A, teal components), we modeled cytoskeletal reorganization, particularly F-actin dynamics through ROCK, mDia, and cofilin and stress fiber accumulation through the cumulative effects of myosin and F-actin. These events were modeled spatially building on the well-mixed model in Sun *et al. (28)*) (Fig. 2).

**Figure 2.**
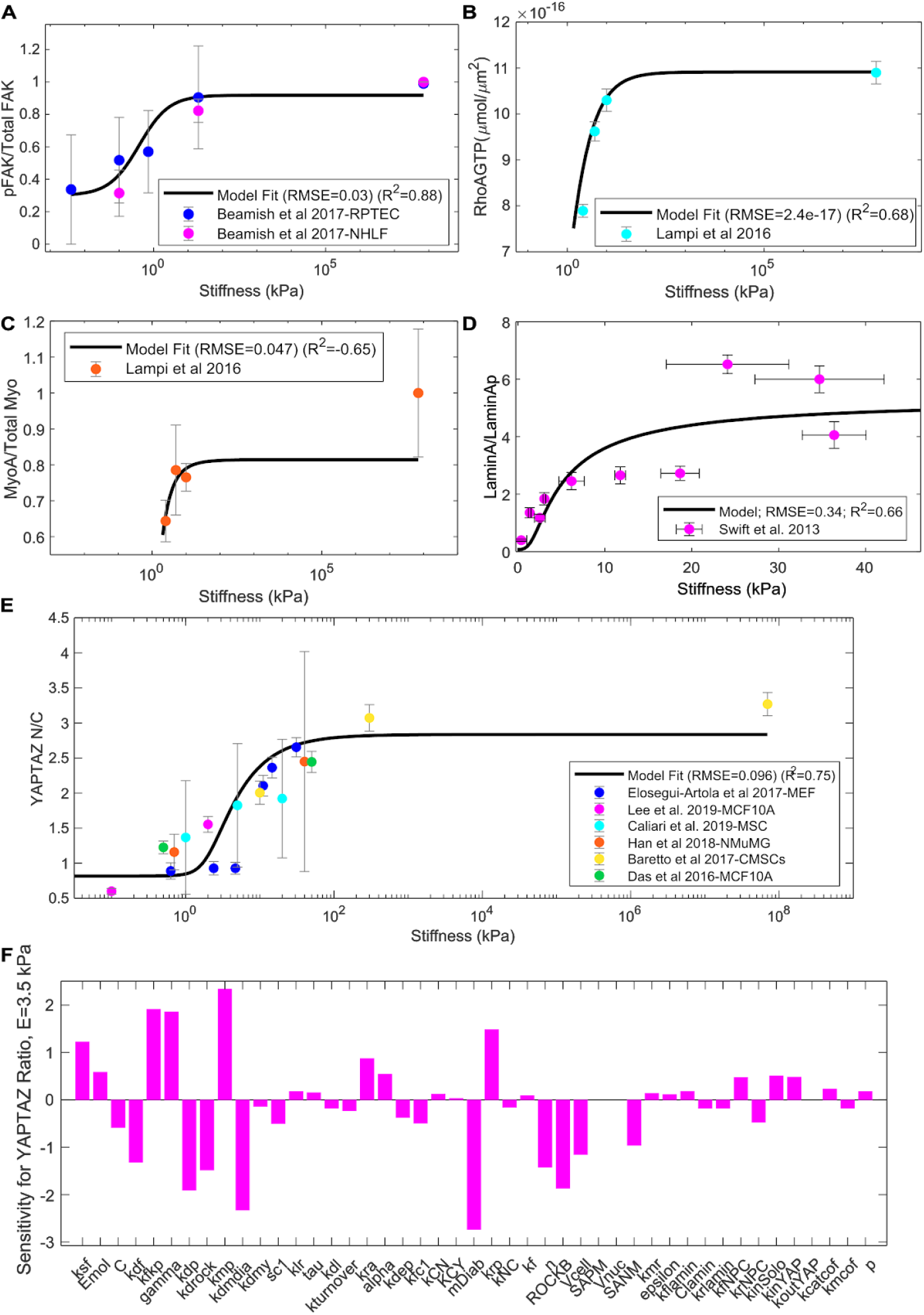
Validation of the compartmental model outcomes against published experiments. (A) The ratio of pFAK to total FAK as a function of stiffness was validated against data from (51) for both fibroblasts (NHLF) and renal proximal tubular epithelial cells (RPTECs). (B) RhoA-GTP activation over stiffness model fit and comparison to experimental data from endothelial cells (34). (C) Ratio of myosin activation over total myosin predicted by our model and a comparison with experimental data from endothelial cells (34). (D) Validation of Lamin A activation as a function of tissue stiffness across various tissues including brain, kidney and bone among others (45). (E) The ratio of nuclear YAP/TAZ to cytosolic YAP/TAZ as predicted by our model and a comparison with experimental data from multiple sources (2, 9, 11, 22, 28, 29). (F) Sensitivity analysis with respect to YAP/TAZ Nuc/Cyto ratio for 3.5 kPa. Additional sensitivity analyses for all of the variables to all of the parameters and all initial conditions can be found in the supplementary material (Figs. S3, S4).

#### 4. RhoA activates mDia

In addition to the activation of ROCK, RhoA also activates mDia. mDia is a formin protein that nucleates actin filaments and accelerates the rate of actin elongation 5-15 fold (28, 38) (Table 2, Rates 7 and 8). Because RhoA-GTP is a membrane bound protein and mDia is in the cytosol, the activation of mDia is modeled as a boundary flux. Inactivation of mDia is modeled as a first-order reaction in the cytosol.

**Table 2:**
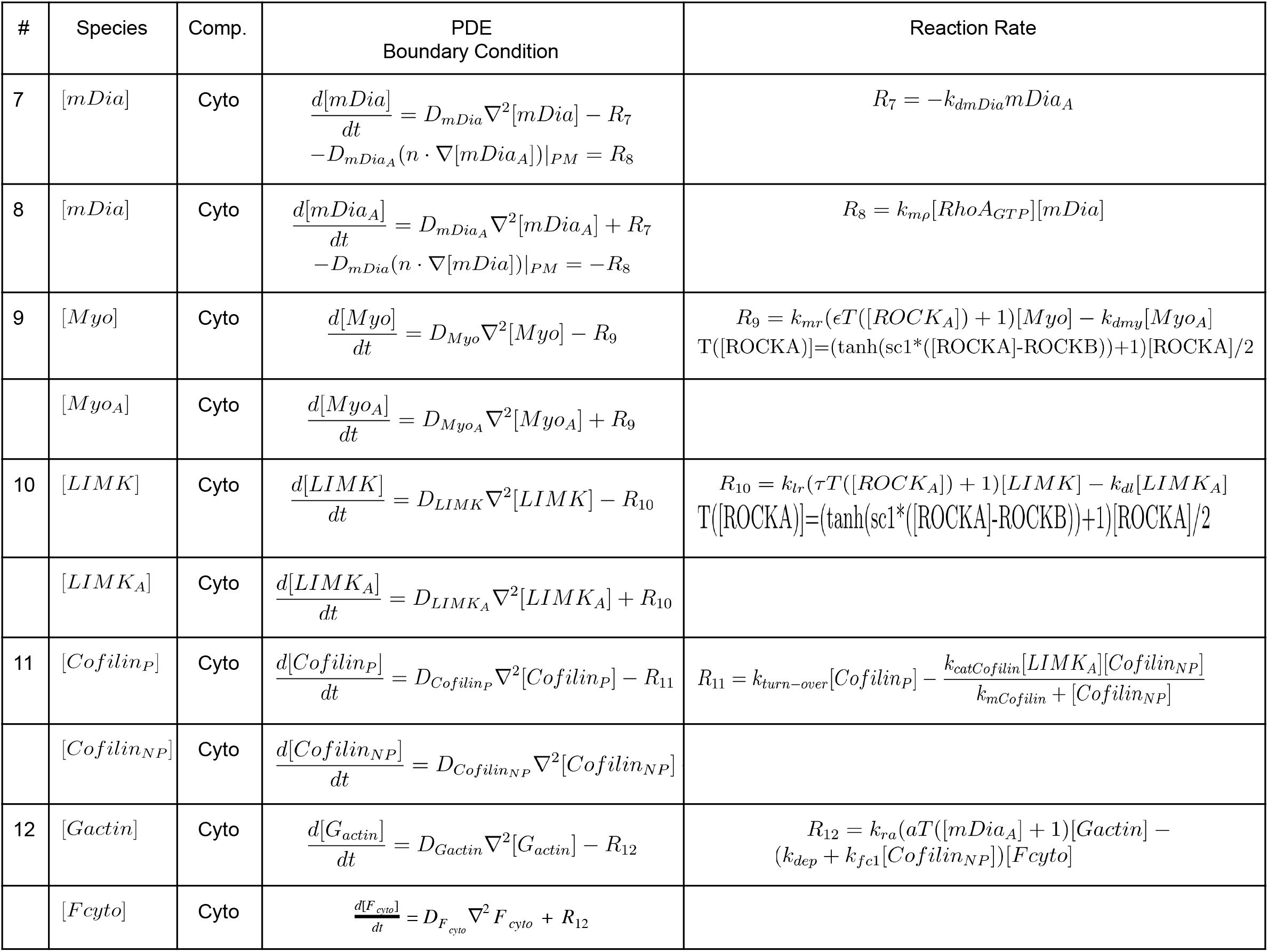
Partial differential equations for the events outlined in Module 2 along with species, locations, spatial equations and reaction rates used in the model. See Table S2 for the definition of the parameters.

| # | Species | Comp. | PDE<br>Boundary Condition | Reaction Rate |
| --- | --- | --- | --- | --- |
| 7 | $[mDia]$ | Cyto | $\frac{d[mDia]}{dt} = D_{mDia} \nabla^2 [mDia] - R_7$ $-D_{mDia_A}(n \cdot \nabla [mDia_A])\vert _{PM} = R_8$ | $R_7 = -k_{dmDia} mDia_A$ |
| 8 | $[mDia_A]$ | Cyto | $\frac{d[mDia_A]}{dt} = D_{mDia_A} \nabla^2 [mDia_A] + R_7$ $-D_{mDia}(n \cdot \nabla [mDia])\vert _{PM} = -R_8$ | $R_8 = k_{mp}[RhoA_{GTP}][mDia]$ |
| 9 | $[Myo]$ | Cyto | $\frac{d[Myo]}{dt} = D_{Myo} \nabla^2 [Myo] - R_9$ | $R_9 = k_{mr}(\epsilon T([ROCK_A]) + 1)[Myo] - k_{dmy}[Myo_A]$ $T([ROCK_A]) = (\tanh(sc1 * ([ROCK_A] - ROCKB)) + 1)[ROCK_A]/2$ |
| | $[Myo_A]$ | Cyto | $\frac{d[Myo_A]}{dt} = D_{Myo_A} \nabla^2 [Myo_A] + R_9$ | |
| 10 | $[LIMK]$ | Cyto | $\frac{d[LIMK]}{dt} = D_{LIMK} \nabla^2 [LIMK] - R_{10}$ | $R_{10} = k_{lr}(\tau T([ROCK_A]) + 1)[LIMK] - k_{dl}[LIMK_A]$ $T([ROCK_A]) = (\tanh(sc1 * ([ROCK_A] - ROCKB)) + 1)[ROCK_A]/2$ |
| | $[LIMK_A]$ | Cyto | $\frac{d[LIMK_A]}{dt} = D_{LIMK_A} \nabla^2 [LIMK_A] + R_{10}$ | |
| 11 | $[Cofilin_P]$ | Cyto | $\frac{d[Cofilin_P]}{dt} = D_{Cofilin_P} \nabla^2 [Cofilin_P] - R_{11}$ | $R_{11} = k_{turn-over}[Cofilin_P] - \frac{k_{catCofilin}[LIMK_A][Cofilin_{NP}]}{k_{mCofilin} + [Cofilin_{NP}]}$ |
| | $[Cofilin_{NP}]$ | Cyto | $\frac{d[Cofilin_{NP}]}{dt} = D_{Cofilin_{NP}} \nabla^2 [Cofilin_{NP}]$ | |
| 12 | $[Gactin]$ | Cyto | $\frac{d[Gactin]}{dt} = D_{Gactin} \nabla^2 [Gactin] - R_{12}$ | $R_{12} = k_{ra}(aT([mDia_A] + 1)[Gactin] - (k_{dep} + k_{fc1}[Cofilin_{NP}])[Fcyto])$ |
| | $[Fcyto]$ | Cyto | $\frac{d[Fcyto]}{dt} = D_{F_{cyto}} \nabla^2 F_{cyto} + R_{12}$ | |

#### 5. ROCK activates Myosin

ROCK promotes myosin activity by phosphorylating myosin light chain kinase, and inhibiting myosin phosphatase, which, in turn, regulates cellular contractility. To capture the nonlinear nature of these interactions, we model the activation of myosin using a hyperbolic tangent function similar to the one described in (28) (Table 2, Rate 9, Fig. S1). Inactivation of myosin is modeled using first order kinetics (Table 2, Rate 9).

#### 6. ROCK activates LIMK

ROCK is one of the key upstream regulators of LIMK and activation of ROCK further activates LIMK (28, 39). To capture the nonlinear interactions of ROCK and LIMK, we use a similar hyperbolic tangent function as described for myosin activation in our model (Table 2, Rate 10) and a first order rate for LIMK inactivation (28, 40, 41).

#### 7. Cofilin inhibits F-actin polymerization

LIMK phosphorylates and inactivates cofilin, an F-actin severing protein (42). These relationships are incorporated into the model as shown in Table 2, Rate 11.

#### 8. F-actin dynamics

The polymerization of F-actin is modeled using two functions. The first function is an mDia*-*dependent polymerization of G-actin to F-actin and the second is a basal polymerization rate of F-actin. We assume that mDia must reach a certain concentration in order to effectively polymerize F-actin and use a hyperbolic tangent function to model this thresholding (Table 2, Rate 12). We do not keep track of the explicit number of barbed ends. Similar to the simplified polymerization model, we include a basal depolymerization rate of F-actin and a cofilin-mediated net depolymerization (28, 40, 41, 43).

### Module 3: YAP/TAZ nuclear translocation

In this module (Fig. 1A, pink components), we designed a set of reactions to reflect the mechanochemical events that affect the nuclear translocation of YAP/TAZ. Nuclear translocation of YAP/TAZ depends on stress fibers, lamin A activation in the nucleus, and nuclear shape and stiffness. By developing the reactions that reflect these events, we complete the set of minimal events that capture the nuclear translocation of YAP/TAZ. Here we introduce two cytosolic species of YAP/TAZ: phosphorylated YAP/TAZ, called *YAP/TAP*_*P*_, and dephosphorylated YAP/TAZ, called *YAP/TAP*_*N*_, which can translocate to the nucleus. *YAP/TAP*_*nuc*_ represents the concentration of YAP/TAZ in the nucleus. We calculate the YAP/TAZ Nuc/Cyto ratio using the relative volumes of each compartment.

#### 9. Stress fibers activate YAP/TAZ dephosphorylation

YAP/TAZ dephosphorylation depends on stress fiber formation from F-actin and myosin (3, 44). In our model, the accumulation of stress fibers in the cell is incorporated as a product of F-actin and myosin (Table 3, Rate 13).

**Table 1:** Partial differential equations for the events outlined in Module 3 along with species, locations, spatial equations and reaction rates used in the model. See Table S2 for the definition of the parameters.

| # | Species | Comp. | PDE<br>Boundary Condition | Reaction Rate |
| --- | --- | --- | --- | --- |
| 13 | $[YAPTAZ_P]$ | Cyt<br>o | $\frac{d[YAPTAZ_P]}{dt} = D_{YAPTAZ_P} \nabla^2 [YAPTAZ_P] - R_{13}$ | $R_{13} = (k_{CN} + k_{CY}[F_{cyto}][Myo_A])[YAPTAZ_P] - k_{NC}[YAPTAZ_N]$ |
| 14 | $[LaminA_p]$ | NM | $\frac{d[LaminA_p]}{dt} = D_{LaminA_p} \nabla^2 [LaminA_p] - R_{14}$ | $R_{14} = k_{fl} \frac{E_{cytosol}}{C_{LaminA} + E_{cytosol}} [LaminA_p] - k_{rl} [LaminA]$<br>$E_{cytosol} = p[F_{cyto}]^{2.6}$ |
| | $[LaminA]$ | NM | $\frac{dLaminA}{dt} = D_{LaminA} \nabla^2 [LaminA] + R_{14}$ | |
| 15 | $[NPC]$ | NM | $\frac{d[NPC]}{dt} = D_{NPC} \nabla^2 [NPC] - R_{15}$ | $R_{15} = k_{fNPC} [LaminA][F_{cyto}][Myo_A][NPC] - k_r [NPC_A]$ |
| | $[NPC_A]$ | NM | $\frac{d[NPC_A]}{dt} = D_{NPC_A} \nabla^2 NPC_A + R_{15}$ | |
| 16 | $[YAPTAZ_N]$ | Cyt<br>o | $\frac{dYAPTAZ_N}{dt} = D_{YAPTAZ_N} \nabla^2 YAPTAZ_N + R_{13}$<br>$-D_{YAPTAZ_N}(n \cdot \nabla YAPTAZ_N)\vert _{NM} = -R_{16}$ | $R_{16} = (k_{in_b} + k_{in} NPC_A) YAPTAZ_N - k_{out} YAPTAZ_{nuc}$ |
| | $[YAPTAZ_{nuc}]$ | Nuc | $\frac{dYAPTAZ_{nuc}}{dt} = D_{YAPTAZ_{nuc}} \nabla^2 YAPTAZ_{nuc}$<br>$-D_{YAPTAZ_{nuc}}(n \cdot \nabla YAPTAZ_{nuc})\vert _{NM} = R_{16}$ | |

#### 10. Lamin A activation due to stiffness

Emerging evidence points to the importance of lamin A in regulating YAP/TAZ nuclear localization (45, 46). Furthermore, Lamin A is sensitive to substrate stiffness. Lamin A molecules form a network of intermediate filaments in the inner nuclear lamina with complex dynamic and mechanosensitive regulation; Lamin A phosphorylation promotes meshwork disassembly and protein turnover. On stiff substrates, Lamin A is dephosphorylated and incorporated into the inner nuclear lamin, while on soft substrates Lamin A is significantly more phosphorylated (45). What is not yet clear is how Lamin A senses substrate stiffness. To simplify the interactions between lamin A and substrate stiffness, we argue that substrate stiffness is read by the upstream events in Modules 1 and 2 and alters cytosolic stiffness as a function of F-actin concentration. We propose a model in which the rate of lamin A dephosphorylation is a function of cytosolic stiffness, where the cytosolic stiffness is a function of F-actin, which, in turn, is responsive to substrate stiffness. To develop this nested model, we first estimated the relationship between F-actin concentration and actin stiffness using data presented in (47). Using this data, we propose that cytosolic stiffness increases as a function of F-actin using a power law with an exponent of 2.6 (see Fig S2 for fit) so that *E*_*cytosol*_ = *p*[*F*_*cyto*_]^2.6^. Then, we propose Lamin A dephosphorylation as a function of cytosolic stiffness, similar to the mechanotransduction model for FAK (Table 3, Rate 14).

#### 11. Stretching of nuclear pores

The interaction between the cytoskeleton and nucleus through lamin A and LINC complexes generates nuclear stress, which then induces stretching of nuclear pores complexes (NPCs), decreasing mechanical resistance and increasing YAP/TAZ nuclear import (17). We model the phenomenon of nuclear stress induced stretching of NPCs using the relationship shown in Table 3 Rate 15 (16).

#### 12. YAP/TAZ nuclear import and export

In this step,YAP/TAZ nuclear import is controlled by basal level YAP/TAZ import and nuclear import due to stretching of nuclear pore complexes and the associated import and export proteins, including the state of Lamin A (6, 8, 17, 30, 48, 49). YAP/TAZ nuclear export is modeled using a simple first-order rate (Table 1, Rate 16) (6, 8, 17, 30, 48, 49). The transport of YAP/TAZ across the nuclear membrane is modeled as a boundary condition for both *YAP/TAZ*_*nuc*_ flux into the nucleus and *YAP/TAZ*_*N*_ transport in the cytoplasm (Table 3, Rate 16).

### Estimation of kinetic parameters using a well-mixed model

The signaling model has a total of 40 kinetic parameters. Twenty-three of these parameters were obtained from the literature, while the remaining 17 were estimated in this work using published experimental data in the literature (Fig. 2). All experimental data were assumed to be at steady state to enable such comparisons, and parameters were estimated using COPASI (50). In order to find each of the estimated parameters, a large range of possible parameters was tested, as shown in Table S3. The finalized list of parameters is given in Table S4 and geometry sizes are given in Table S5. Initial conditions were estimated based on the total amount of the relevant protein from the literature (see Tables S7 and S8). Local sensitivity analysis was conducted to identify the effect of different initial conditions and kinetic parameters on model response (Fig. S3, S4).

### Validation of the well-mixed model with published experimental results

We validated the simulation output from the model against experiments in the published literature (Fig. 2). We focused on the relationship between stiffness of the substrate and the activation of key species including FAK, RhoA, myosin, Lamin A, and YAP/TAZ Nuc/Cyto ratio. The activation of FAK in response to stiffness, denoted by pFAK, was in good alignment between model and experiment (RMSE=0.03; R^2^=0.88) (Fig. 2A). Good agreement was also observed between RhoA-GTP membrane species and experimental results from Lampi *et al.* (RMSE=2.4 × 10^−17^; R^2^=0.68) (34) (Fig. 2B). Our model also showed a good agreement between experiments and simulations for Lamin A activation as a function of substrate stiffness (Fig. 2D). Thus, our proposed multiscale model of substrate stiffness impacting cytosolic stiffness, which in turn alters Lamin A activation was able to capture the experimentally observed relationship between substrate stiffness and Lamin A (RMSE=0.34; R^2^=0.66). On the other hand, there was good agreement with the model for myosin activation for low stiffness, but poor agreement for myosin activation at high stiffness such as glass (RMSE=0.047, R^2^=−0.65; the negative R^2^ value indicates that mean value would have been a good fit) (Fig. 2C). This is probably because of missing biochemical interactions in the model. However, we prioritized the prediction of the myosin activation for low stiffnesses due to the importance of YAP/TAZ nuclear localization at physiologic stiffnesses. We also conducted parametric sensitivity analysis for three different values of stiffnesses, 0.1 kPa, 3.5 kPa and 5.7 kPa.

We conducted sensitivity analyses for all the parameters, including the kinetic parameters, compartment sizes, and the initial concentrations of the different species. This analysis shows that, in our model, YAP/TAZ Nuc/Cyto ratio is sensitive to the parameters including FAK activation due to stiffness including *k*_*s,f*_, *C*, and *E* (Fig. 2E). At very low (0.1 kPa) and high (5.7 kPa) stiffnesses, YAP/TAZ Nuc/Cyto ratio is less sensitive to the stiffness related variables (Fig. S2A). At high stiffnesses, YAP/TAZ is very sensitive to other parameters of the model including those associated with RhoA-GTP activation, ROCK activation, and mDia activation (Fig. 2F). This sensitivity is due in part to the fact that the YAP/TAZ Nuc/Cyto does not change significantly at high stiffnesses, so any small change in kinetic parameters leads to larger downstream effects. Further studies could validate the RhoA-GTP, mDia, and ROCK activity. YAP/TAZ Nuc/Cyto ratio as well as other variables of the model are sensitive to the initial concentration of FAK, RhoA, mDia and ROCK. Additional sensitivity analysis for the initial conditions and parameters for all variables of the model can be found in Fig. S3 and S4. Having constrained the relevant parameters, we next developed a spatial model of YAP/TAZ nuclear translocation.

### Translation from well-mixed to spatial models

All simulations are conducted in 3D (Fig. 3–7) using the Virtual Cell modeling suite (52, 53) and results from the simulations are shown as a 2D cross section for ease of interpretation. The full 3D geometry can be visualized by rotating around the vertical axis of symmetry. We constructed a cellular geometry to represent a cell that has spread on a flat 2D surface. As a cell spreads, the length of the cell tapers so that the tip of the cell becomes thin (54). On stiff substrates, the cell tends to spread and become elongated. On soft substrates, the cell remains rounded (8). Diffusion coefficients are given in Table S6. We parameterized a series of geometries in Virtual Cell to test the effect of different cell and nuclear shapes (Tables S9, S10).

**Figure 3.**
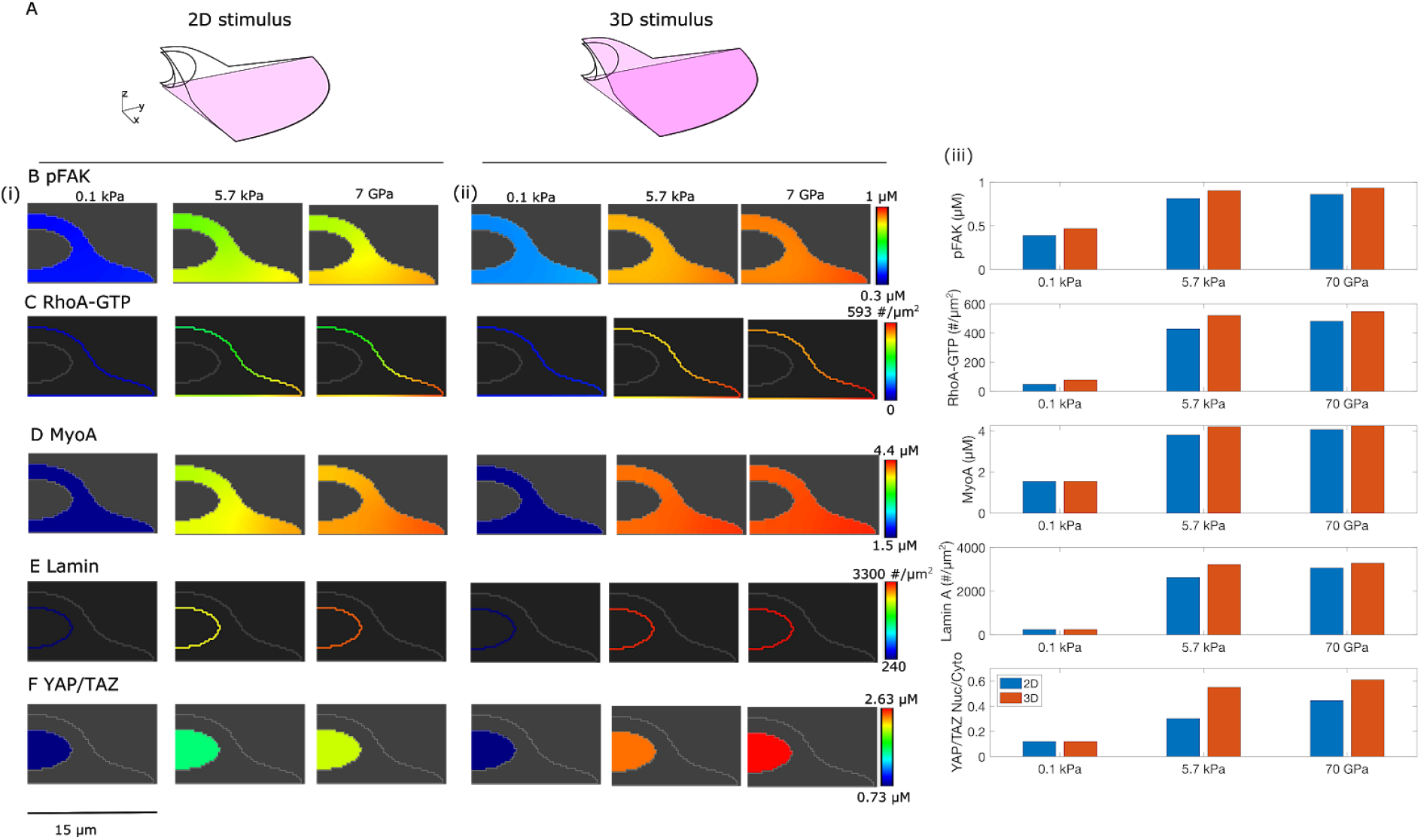
Effect of substrate dimensionality and substrate stiffness on YAP/TAZ nuclear translocation. (A) Schematic showing the model setup for substrate dimensionality presentation. (B) Steady state profiles at 4000 s for (A) pFAK, (C) RhoA-GTP, (D) MyoA, (E) Lamin A, and (F) YAP/TAZ nuclear activation. In each row, (i) shows the effect of stiffness when the stimulus is presented in 2D, (ii) shows the effect of stiffness when the stimulus is presented in 3D, and (iii) shows a direct comparison between the 2D and 3D stimulus of the same values in a bar graph. For YAP/TAZ the nuclear to cytosolic ratio is shown in panel F(iii).

### 2D versus 3D substrate presentation

In 2D, cells are able to interact with the substrate only on the basal side, however in 3D, the substrate surrounds the cell on all sides. To simulate 2D and 3D culture conditions, we varied the localization of the boundary condition of stiffness. Substrate stiffness was implemented as a boundary condition on the plasma membrane for all the simulations. For simulations of cells on 2D surfaces, we modeled the substrate contact area on the basal, flat side of the cell where the cell would touch the surface of the coverslip or gel in 2D (as shown in pink in Fig. 3A, left). For simulations of cells in 3D substrates, we localize stiffness activation to the entire surface of the plasma membrane (as shown in pink in Fig. 3A, right). Although the cell interacts with the matrix at discrete locations, i.e. focal adhesions, we model these activation areas as continuous for ease of computation, thereby restricting the number of free parameters.

### Simulation methods

The system of partial differential equations resulting from the reaction diffusion system was solved numerically using the Virtual Cell suite (52, 53). We used a fully-implicit finite volume, regular grid (variable time step) solver with an absolute and relative error tolerance of 1×10^−9^ and 1 1×10^−7^ respectively and the maximum time step as 0.1 s. Simulations were run for a total of 4000 seconds to obtain steady state. The mesh size for each geometry is approximately 0.26 μm and the mesh is uniform in X, Y, and Z. All Virtual Cell modeling files will be made publicly available upon publication of the manuscript.

## RESULTS

Using the model developed above, we investigated how the shape of the cell and nucleus, as well as the localization of the stiffness signal could affect YAP/TAZ nuclear localization. To analyze these coupled effects, we have organized our simulations as follows. First, we investigate the effect of substrate dimensionality and substrate stiffness on YAP/TAZ nuclear localization (Figs. 3 and 4). Next, we investigate the effect of cell shape changes that maintain constant substrate activation area across different substrate dimensionalities and their impact on perceived stiffness in terms of YAP/TAZ Nuc/Cyto ratio (Figs. 5 and 6). Finally, we change the shape of the nucleus (Fig. 7). We describe the predictions of these simulations in detail below.

**Figure 4.**
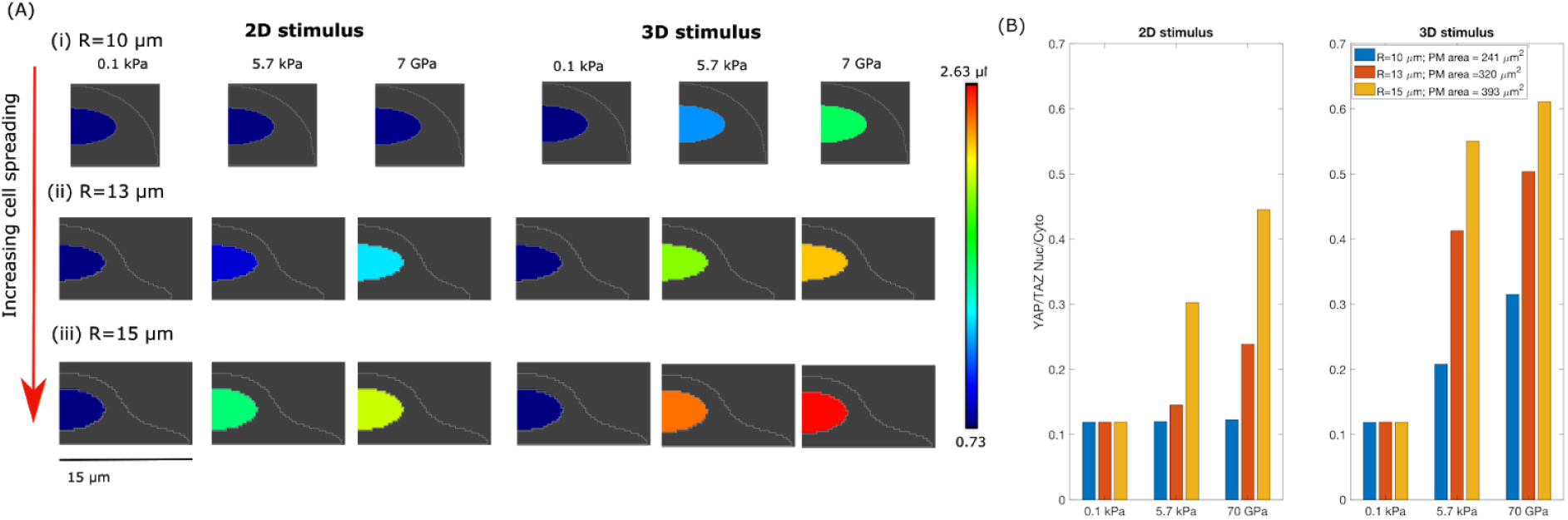
YAP/TAZ nuclear localization depends on cell shape, substrate stiffness, and substrate dimensionality. (*A*) YAP/TAZ nuclear localization at 4000s for 3 different geometries representing increasing cell elongation and three different substrate stiffnesses (i) R=10 μm, plasma membrane area=241 μm^2^, (ii) R=13 μm, plasma membrane area=320 μm^2^ and (iii) R=15 μm, plasma membrane=393 μm^2^. (B) YAP/TAZ Nuc/cyto ratio for the three different cell sizes for three different stiffness values (physiologic stiffness ~0.1 kPa in blue, tumor stiffness ~5.7 kPa in red, and glass ~70 GPa in yellow).

**Figure 5:**
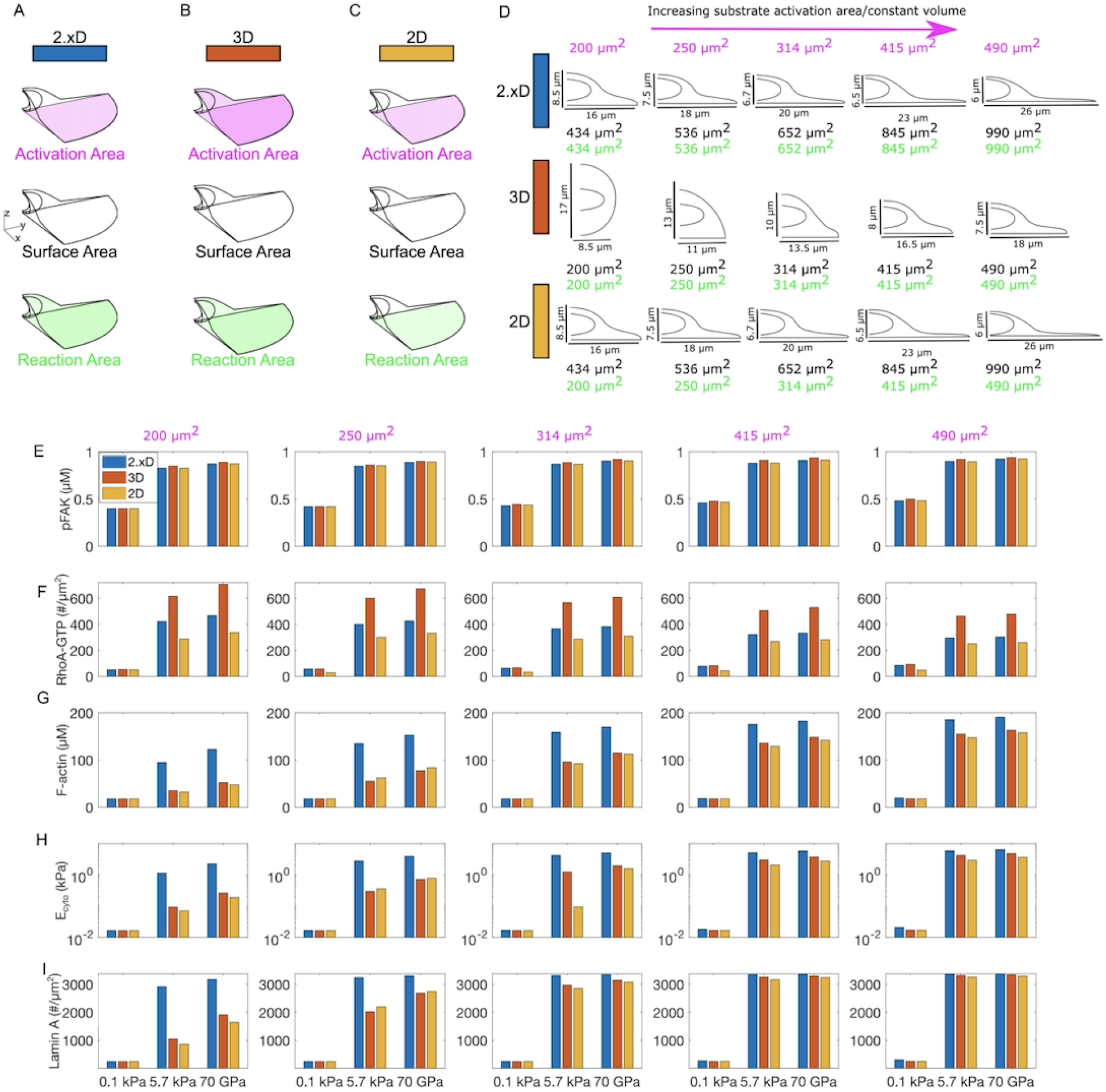
Differences in substrate presentation and membrane reaction area alter responses to 2D and 3D stimulus. (A) 2.xD substrate presentation where the substrate is present only on the bottom surface but the entire plasma membrane is available for membrane reactions. (B) 3D substrate presentation where the substrate activation and the membrane reactions occur on the entire plasma membrane. (C) 2D substrate presentation with the same area used for substrate presentation and for membrane reactions. (D) Increasing substrate activation area while keeping cytosolic volume constant forces cell shape changes dependent on stimulus presentation geometry. (E) pFAK, (F) RhoA-GTP, (G) F-actin, (H) Ecyto, and (I) Lamin A as a function of substrate presentation, substrate stiffness, and activation area.

**Figure 6.**
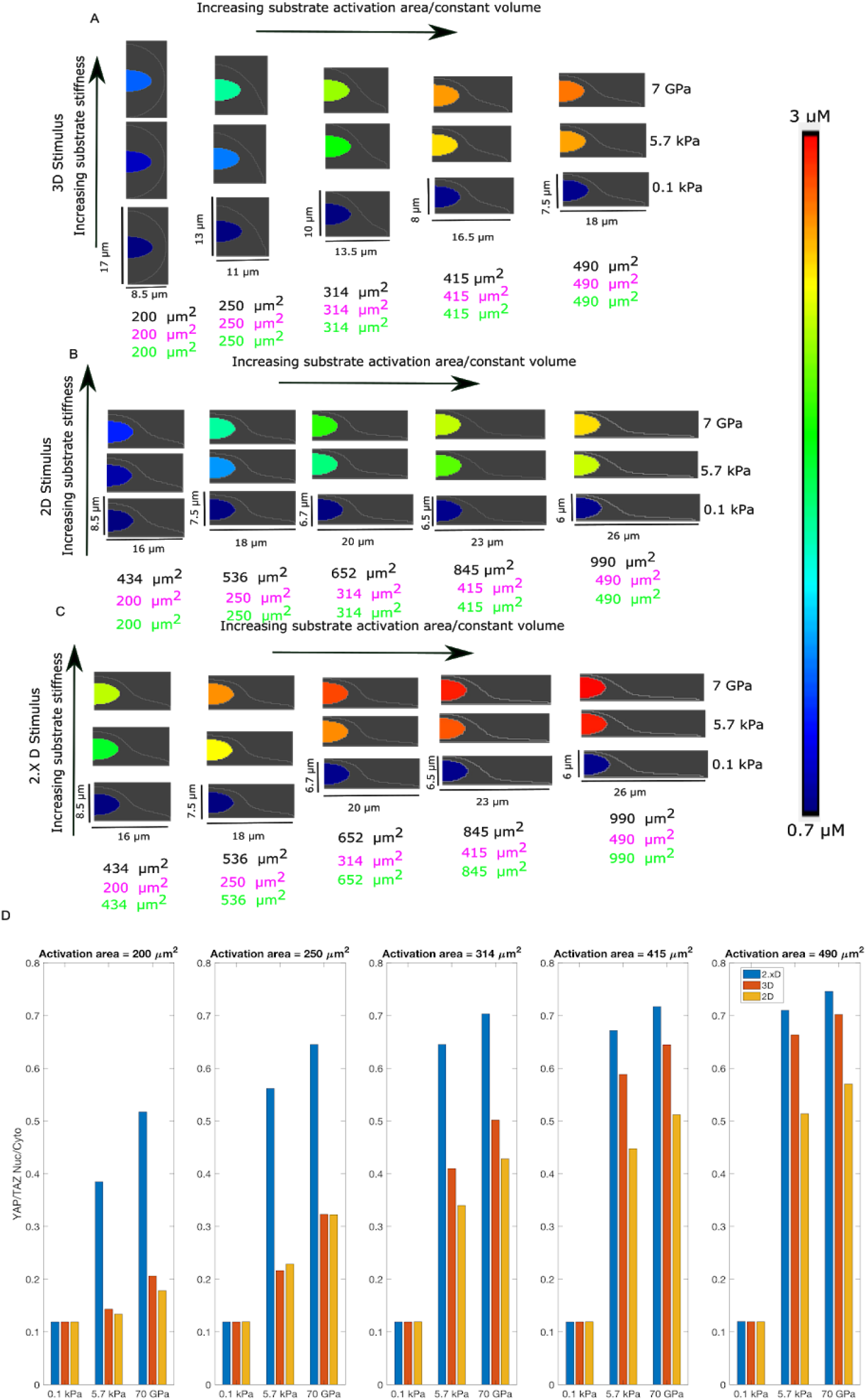
Increasing substrate activation area changes YAP/TAZ Nuc/Cyto ratio as a function of substrate dimensionality. Nuclear YAP/TAZ for different activation areas for (A) 3D stimulus, (B) 2D stimulus, and (C) 2.X D dimensions. The black, pink, and green surface areas refer to the total plasma membrane surface area, the surface area activated by the substrate stiffness, and the surface area available for membrane reactions, respectively, as detailed in Fig. 5D. (D) YAP/TAZ Nuc/Cyto ratio for different activation areas and different stimuli.

**Figure 7.**
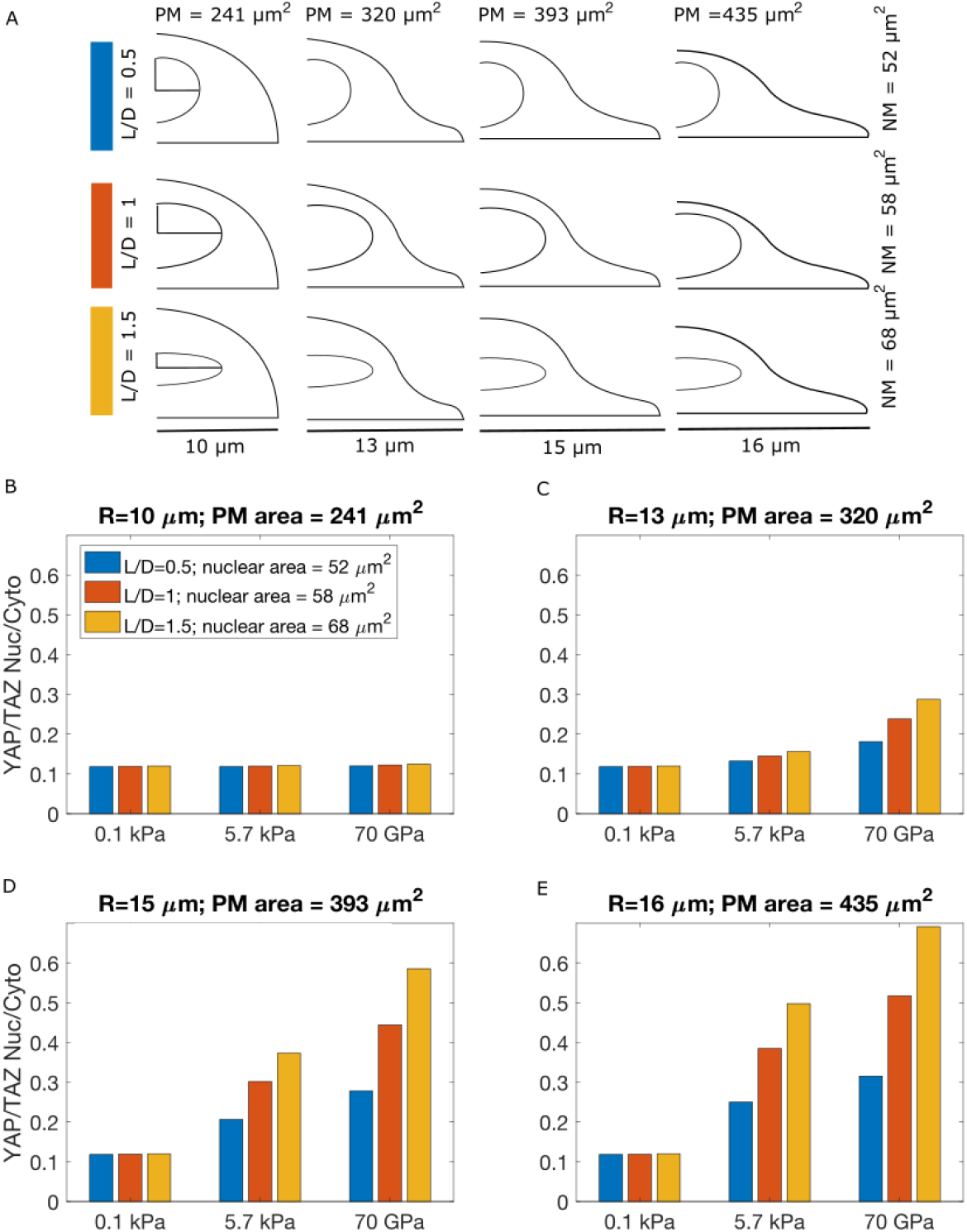
YAP/TAZ Nuc/Cyto is influenced by nuclear shape. YAP/TAZ Nuc/Cyto ratio for four different cell sizes and three different nuclear shapes. (A) The nuclear shapes were varied by changing the length/diameter ratio, while conserving the nuclear volume. (B) R=10 μm, plasma membrane area = 241μm^2^, (C) R=13 μm, plasma membrane area = 320 μm^2^, (D) R=15 μm, plasma membrane area = 393 μm^2^, and (E) R=16 μm, plasma membrane area = 435 μm^2^. Blue bars L/D=0.5, nuclear membrane area = 52 μm^2^; Red bars L/D=1, nuclear membrane area = 58 μm^2^; yellow bars L/D=1.5, nuclear membrane area = 68 μm^2^.

**Figure 8.**
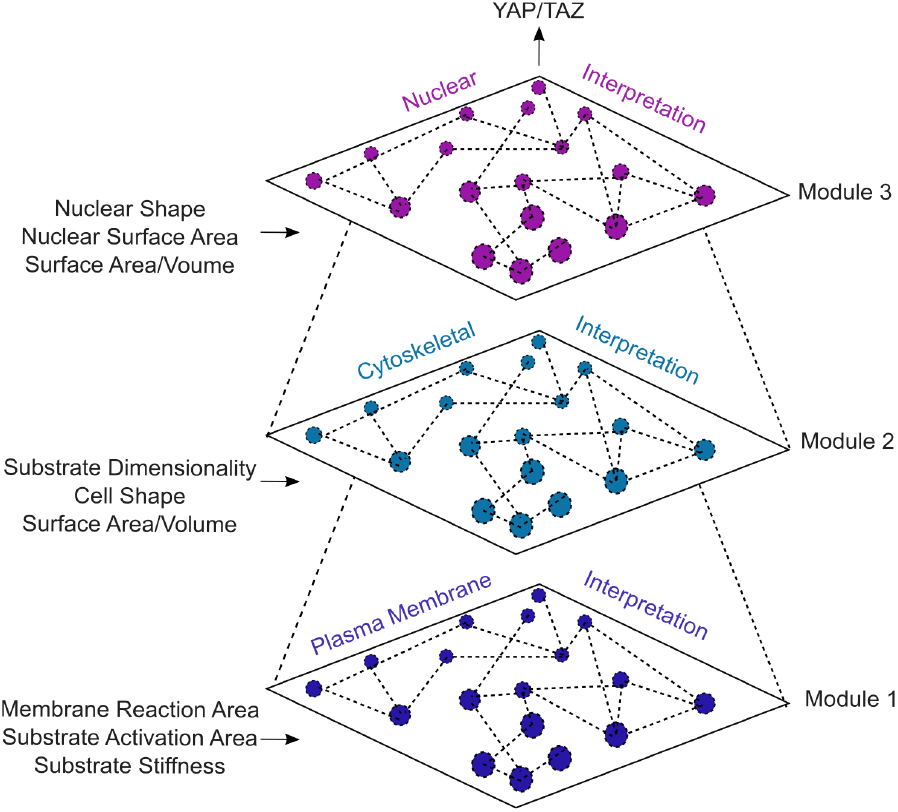
YAP/TAZ nuclear localization outcomes rely on multiplexed signals integrated through the cellular transfer function. In addition to substrate stiffness, other physical cues must be interpreted by the cell through the plasma membrane, cytoskeleton, and nucleus via layers of information transfer. Such cues include the dimensionality of the substrate, i.e. 2D versus 3D, cell shape restrictions imposed by the ECM, and plasma membrane and nuclear membrane surface area.

### YAP/TAZ nuclear localization depends on substrate stiffness and substrate dimensionality

Experimental evidence in the literature demonstrates that FAK phosphorylation, RhoA, MyoA, and Lamin A are all functions of substrate stiffness (34, 45, 51). Having demonstrated that the well-mixed model does indeed capture these experimental trends (Fig. 2), we next asked how this complex relationship is affected by cell shape and substrate dimensionality. To answer this question, we conducted simulations in three different geometries representing different spread states, 30 *μm* diameter cell with a thin lamellipodia, 20 *μm* with no thin lamellipodium, and an intermediary cell shape with 26 *μm* diameter (Ditlev et al. 2009; Caliari et al. 2016), and three different stiffness values representing normal tissue ~0.1 kPa, tumorigenic stiffness, 5.7 kPa (55) and glass 70 GPa (56) (Fig. 3). The boundary conditions for the 2D versus 3D substrate presentation are shown in (Fig. 3A). We first looked at how components in Module 1 respond to changes in stiffness, shape, and dimensionality. For a given shape, we found that substrate stiffness does indeed increase the amount of pFAK in a cell and that this effect is somewhat increased in 3D compared to 2D (Fig. 3B). The differences in pFAK for 2D versus 3D substrate presentation are small and the stiffness effects are dominant. Downstream of FAK, RhoA-GTP (Fig. 3C), MyoA (Fig. 3D) both show strong stiffness dependence, indicating that the effect of stiffness is propagated to the downstream membrane and cytosolic components in Module 1. We also note that RhoA-GTP, which is membrane bound, demonstrates a strong cell shape dependent gradient along the membrane, consistent with our previous observations (21). The effect of 2D versus 3D presentation is not a dominant feature for these components in Module 1, while stiffness dominates the response.

Downstream of FAK and RhoA-GTP, we looked at the different components in Modules 1 and 2 (Fig. S6) and noted that all components showed a stiffness and dimensionality proportional response (Fig. S6A-E). Interestingly, the cytosolic stiffness shows a dramatic change in response to increasing substrate stiffness for a given substrate dimensionality (Fig. S6F). The power law dependence of cytosolic stiffness on actin concentration means that small changes in F-actin are magnified dramatically in the cytosolic stiffness. This is evident then in Lamin A activation (Module 3) (Fig. 3E). We observed that Lamin A activation integrates volumetric biochemical and mechanical signals onto the nuclear membrane surface and shows dependence on both substrate stiffness and substrate dimensionality. For a given substrate stiffness, Lamin A activation is slightly higher in 3D than in 2D. However, for a given substrate dimensionality, Lamin A activation increases with stiffness (Fig. 3E). Finally, the small changes in Modules 1 and 2 integrate into larger changes in Module 3 to result in both stiffness and dimensionality dependent YAP/TAZ nuclear activation (plotted as YAP/TAZ nuclear concentration in Fig. 3F(i) and (ii), and as YAP/TAZ nuclear to cytosolic ratio in (iii)). Interestingly, at the lower end of the substrate stiffness spectrum modeled here, YAP/TAZ is more sensitive to stiffness changes than dimensionality changes (compare 0.1 kPa to 5.7 kPa in Fig. 3B iii). However, at the higher end of the substrate stiffness spectrum, YAP/TAZ is more sensitive to dimensionality changes than stiffness changes (compare 5.7 kPa to 70 GPa in Fig. 3B iii). In summary, each upstream component in the pathway responds strongly to stiffness changes, with little difference between the 2D versus 3D response. Dimensionality changes are sensed downstream, when YAP/TAZ integrates signals directly from the cytoskeleton and indirectly from the effect of cytosolic stiffness on the nucleus.

### YAP/TAZ nuclear localization integrates cell shape and substrate presentation information

Depending on the culture dimensions, cell shape can vary widely from elongated shapes in 2D culture to rounded shapes in 3D culture (57, 58). Given the mixed conclusions in the literature on the role of culture dimensions on YAP/TAZ nuclear localization (Fig. 1C), we next asked if we could use the 3D spatial model to dissect the effects of cell shape, substrate stiffness, and substrate dimensionality on YAP/TAZ nuclear localization. To investigate how these different effects could play a role, we systematically varied cell shape while conserving cell volume and nuclear dimensions and used 2D or 3D substrate stimulus for the same three stiffnesses as before (Fig. 4). We note that increasing cell elongation is the equivalent to increasing the surface area of the plasma membrane for activation by substrate stiffness and reaction of plasma membrane associated components, i.e. an increase in the surface area to volume ratio of the cell. For a given cell shape, all components in the signaling networks in Modules 1 and 2 showed the same stiffness dependent response as expected (Fig. S7) and in all cases the 3D responses were slightly larger than the 2D stimulus response. Low substrate stiffness values (0.1 kPa) did not elicit any noticeable shape dependent effects, while the intermediate (5.7 kPa) and high (70 GPa) stiffnesses showed dependence on cell shape. Spread cells, with a larger radius had a larger response for various signaling components for a given value of stiffness (Fig. S7A-G, with the exception of Fig S7B, Rho-GTP) and this effect was magnified in the readout of cytosolic stiffness (Fig S7H) and Lamin A activation (Fig S7I). As a result of these upstream effects, we found that for a given cell shape, increasing stiffness increased the YAP/TAZ Nuc/Cyto ratio and this response was always amplified in 3D versus 2D by Module 3 of the pathway (Fig. 4A, B), consistent with what we observed in Fig. 3. We also found that increased cell elongation, with an effectively larger plasma membrane surface area that could result from cell spreading or other effects of 2D culture, resulted in higher YAP/TAZ Nuc/Cyto ratio when compared to rounded cells. Thus, for a given stiffness stimulus, increased cell elongation leads to increased YAP/TAZ Nuc/Cyto ratio and this effect is more pronounced in 3D substrate presentation rather than 2D substrate presentation. Thus, cell elongation amplifies stiffness sensing and dimensionality sensing by the YAP/TAZ pathway.

Thus far, simulations from the model show that maintaining the same cell shape but changing the activation area so that stiffness is applied on the entire cell boundary (simulating 3D culture) increases YAP/TAZ nuclear localization relative to stimulus on the basal side only (simulating 2D culture) (Fig. 3F, 4). This result is not very surprising, since the total surface activation area, and thus the total signal, increases from 2D to 3D conditions. However, experimental data shows that lower YAP/TAZ nuclear localization can sometimes occur in 3D compared to 2D culture in response to stiffness increases (29, 30). This reveals that a simple change in substrate dimensionality alone is insufficient to explain YAP/TAZ signaling differences observed experimentally. We note that there are major differences between 2D and 3D substrate presentation for a given cell shape. Currently, the literature in the field suggests that in 2D culture, adhesion signaling originates from the basal side of the cell, whereas in 3D culture, adhesion signaling is localized across the entire cell surface (14, 59–61). Our model captures this effect since the activation area for FAK, which is an immediate downstream readout of substrate stiffness is different in each case because the surface area of the membrane that is exposed to the substrate in these two different culture conditions is different (Fig. 3A). However, the surface area available for the membrane associated reactions downstream of FAK activation is the same in 2D and 3D and is the total surface area of the plasma membrane. For example, RhoA-GTP, which is membrane-bound, shows a different response from other components in the case of increasing cell elongation due to the differences in membrane activation downstream of FAK (Fig. S7B). These results suggest that the differences in surface presentation of the stimulus to upstream components versus surface area available for all the downstream membrane reactions can lead to confounding results for nuclear localization of YAP/TAZ. We next explore this concept in our model.

### Differences in surface-to-volume ratio for substrate activation versus surface-to-volume ratio for plasma membrane reactions dictate the trends of YAP/TAZ nuclear translocation

To investigate the role played by substrate activation area versus the total plasma membrane area in signal integration and amplification, we conducted the following simulations. We classified our geometries into three categories: (i) “2.xD” stimulus is defined by substrate activation of FAK on the basal side of the cell with membrane reactions occurring across the entire plasma membrane area (Fig. 5A) (see supplementary material for details of boundary condition implementation), (ii) “3D” stimulus is defined by substrate activation of FAK on the entire plasma membrane area as well as membrane reactions occurring across the entire plasma membrane surface area (Fig. 5B), and (iii) “2D” stimulus is defined by substrate activation of FAK and all membrane reactions restricted to the basal side of the cell (Fig. 5C). All geometries had the same cytosolic volume and nuclear dimensions, as a result of which we could draw direct comparisons between the different stimuli presentation and interpretation effects. We conducted these simulations for five different values of substrate activation areas (Fig. 5D). Holding cytosolic volume constant while increasing substrate activation area and varying stimulus presentation geometry resulted in cell shape changes as shown in Fig. 5D.

We found that for a given substrate activation area, pFAK was constant for the three different substrate and membrane area combinations and only depended on substrate stiffness (Fig. 5E). pFAK increased with substrate stiffness, consistent with our previous observations and also increased with substrate area. Thus, the first interpretation of substrate stiffness by the cell appears to be independent of substrate dimensionality. Looking at RhoA-GTP, which uses both the pFAK signal and also the membrane surface area, we found that the surface density of RhoA-GTP was higher in 3D than in 2D where the membrane area and substrate activation area were held constant with our previous results (Fig. 5F). However, the 2D case where the substrate activation area was in the bottom only but the entire plasma membrane area was available for membrane reactions showed Rho-GTP density that was in between the 2D response and the 3D response (Fig. 5F, blue bars). We note that this response was between 2D and 3D, which supports our naming of this situation as “2.xD”. Downstream of RhoA-GTP, we found that the 2.xD case had higher F-actin than 3D or 2D stimulus (Fig. 5G). This can be understood by thinking about the two surface-to-volume ratios at play here. For FAK, the surface-to-volume ratio is the surface area of the membrane that senses the substrate stiffness to the cytosolic volume (which is the same across all conditions). For RhoA, the surface area is the total plasma membrane surface area in 2.xD, which is higher than the substrate presentation area. Therefore, in 2.xD, components in Module 1, respond to a surface-to-volume ratio of 2D substrate presentation, while components in Module 2 and 3, respond to a surface-to-volume ratio of 3D substrate presentation (where the substrate sensing area is the same as the total plasma membrane area). This change in the trend of substrate dimensionality response of F-actin is further amplified in the cytosolic stiffness because of their power law dependence (Fig. 5H). As a result, Lamin A activation on the nuclear membrane is larger for 2.xD than for 3D or 2D (Fig. 5I).

All of these relationships hold true for a given substrate activation area. However, the differences between 2D, 2.xD, and 3D are much larger for smaller activation areas when compared to larger activation areas. For larger activation areas, the differences in the response of various signaling components are smaller for different substrate dimensionalities. This is because as the substrate activation area increases and the cytosolic volume is held constant, all substrate presentation geometries elicit a more elongated cell shape where high surface area-to-volume effects dominate (Fig. 5D). In such shapes, the cytoskeleton module is impacted by the closer proximity of membrane reaction.

Finally, we looked at the effects of the differences in surface activation area and membrane reaction area on YAP/TAZ Nuc/Cyto ratio (Fig. 6). We found that at low stiffness values in 2.xD, 2D, and 3D, increasing surface activation area did not elicit any differences in YAP/TAZ nuclear localization (Fig. 6A-C). At higher stiffness values, YAP/TAZ responded strongly to increasing surface activation area, with the strongest response occurring in 2.xD. As the activation area increased, the 2.xD conditions consistently had higher YAP/TAZ nuclear to cytosolic values for higher stiffness values than 3D or 2D (Fig 6C, D). These results suggest that each module in this mechanochemical network integrates not only the biophysical cues of substrate stiffness but also the geometric information of the cell, particularly the plasma membrane surface area and cytosolic volume to modulate the YAP/TAZ Nuc/Cyto ratio.

### Nuclear shape influences YAP/TAZ nuclear localization

Finally, we explored if alterations to nuclear shape would alter YAP/TAZ Nuc/Cyto ratio. In all our previous simulations, we maintained the nuclear shape (both nuclear membrane area and nuclear volume) to be the same for all simulations. However, it has been reported that nuclear shape can change with cell shape (62, 63). Furthermore, nuclear flattening may drive stretching of the nuclear pore, which can stimulate an increase in YAP/TAZ nuclear import (17). It is thought that nuclear flattening as a result of the stress fibers interaction with LINC complexes has been shown to increase YAP/TAZ Nuc/Cyto in 2D, whereas 3D interaction between stress fibers and nucleus is reduced (16). Therefore, we investigated how nuclear shape alone can alter the YAP/TAZ Nuc/Cyto ratio when a 2.xD stimulus is presented. To test the role of nuclear shape and its interaction with cell shape on YAP/TAZ Nuc/Cyto ratio, we set up the following simulations in 2.xD: we used four different cell shapes ranging from rounded to elongated (R=10 μm to R= 16 μm) and for each cell shape, varied the nuclear shape by changing the length to diameter ratio (L/D ratio) such that the nuclear volume was conserved but the nuclear membrane surface area was changed (Fig. 7A). Again, we observed an interesting dependence of cell shape and nuclear shape on YAP/TAZ nuc/cyto ratio (Fig. 7B-E). First, for the small, rounded cell, there was no strong dependence of YAP/TAZ nuc/cyto ratio on nuclear shape or substrate stiffness (Fig. 7B). Second, for low stiffness, there was no cell or nuclear shape dependent response (Figs. 7B-E, 0.1 kPa). For tumor stiffness values, as the cell shape changes from rounded to elongated (through increasing values of R), we found that cells with elongated nuclei (i.e. larger nuclear membrane area) showed larger YAP/TAZ nuc/cyto ratio. For example, in the cell with R=16 μm, at 5.7 kPa, there is approximately a 2-fold increase in YAP/TAZ Nuc/Cyto ratio when L/D changes from 0.5 to 1.5 (corresponding to an increase in the nuclear surface area). At the stiffness values of 70 GPa, this fold change is greater than 2. Thus, our model predicts that in addition to the substrate activation area and the plasma membrane surface area, the nuclear membrane area also plays an important role in multiplexing the transfer of information from the substrate to the nucleus. Therefore, interpretation of substrate stiffness is fundamentally dependent on cell shape at both the cytosolic and the nuclear level.

## Discussion

In this study, we have constructed a 3D spatial model of YAP/TAZ nuclear translocation in response to substrate stiffness to identify the transfer function(s) that govern this fundamental mechanotransduction pathway. Using a systematic approach to vary substrate stiffness and geometry, we found that upstream components in this transfer function respond to stiffness changes while dimensionality changes are sensed downstream, when YAP/TAZ integrates signals directly from the cytoskeleton and indirectly from the effect of cytosolic stiffness on the nucleus. This highlights a need to investigate how other mechanotransduction circuits like MRTF-SRF, which directly regulate cytoskeleton components, impinge on YAP/TAZ dimensionality sensing. Indeed, experimental evidence is emerging that demonstrates the co-dependence of MRTF-SRF and YAP/TAZ pathways(64).

We also systematically varied cell shape and nuclear shape in our model, since shape is an important cellular property associated with YAP/TAZ signaling (65), but often changes concomitantly with culture dimensionality ECM stiffness, making these features difficult to decouple experimentally. This investigation revealed that how a cell senses stiffness is very tightly regulated by cell shape through different surface-to-volume relationships. Interestingly, we found that low stiffness, corresponding to physiological tissue stiffness was fundamentally a robust state for YAP/TAZ nuclear translocation and changes in cell shape, substrate dimensionality, and nuclear shape did not affect this outcome. Separately, we note that rounded cells were fairly robust to changes in stiffness and substrate dimensionality. Thus, high substrate stiffness alone is not enough to generate a response in YAP/TAZ nuclear translocation, nor is 3D substrate dimensionality. Rather, cell shape and nuclear shape are major determinants of the YAP/TAZ response to substrate stiffness and dimensionality. Thus, experimental evidence that conflicts on the role of substrate stiffness and dimensionality (Fig. 1B, C) would be better compared by considering cell and nuclear shape along with the plasma membrane area in contact with the substrate. Our model predicts that cells in 3D will have less YAP/TAZ activation than cells in 2D when the substrate activation area is small and cells in a 3D substrate must be more rounded than the cells modeled as being in contact with a 2D substrate in order to maintain constant cell volume and activation area. Importantly, these findings also highlight that the multiple mechanisms regulating cell and nuclear shape, such as outside-in changes imposed by the extracellular matrix or inside-out changes directed by gene expression, will determine how stiffness and dimensionality are sensed by the cell, making the landscape of YAP/TAZ nuclear translocation quite complex. For example, nuclear shape is dysregulated in cancer (66, 67) and laminopathies (68). In cells with lamin mutations derived from laminopathies, YAP/TAZ actually tends to be constitutively nuclear. Experimentally, YAP/TAZ nuclear import increases with either nuclear pressure or substrate stiffness, where nuclear pores diameter increases with increasing substrate stiffness (9). Thus, one possibility is that constitutive YAP/TAZ nuclear import occurs through dysregulation of nuclear shape and nuclear pores that can occur in laminopathies (17, 69). Given the multitude of ways in which cell shape is regulated, additional variables may need to be considered in a comprehensive model in order to fully explain differences in YAP/TAZ nuclear localization due to dimensionality and stiffness.

In addition to cell shape, nuclear shape, and substrate activation area, our model suggests that the cell’s surface area that is available for membrane reactions is another dominant factor in YAP/TAZ mechanotransduction. Experimentally, it has been observed that increasing cell surface area corresponds to increasing YAP/TAZ Nuc/Cyto (70, 71). While membrane mechanotransduction is typically attributed to proteins, like integrins and ion channels in the plasma membrane and lamins associated with the nuclear membrane, cell surface area regulation depends on phospholipid and glycocalyx polymer trafficking (72). Most cells use membrane trafficking to continually rework their plasma membrane through ruffles, folds, endo- and exocytosis, and it has been suggested that cells sense and regulate their volume and surface area through plasma membrane tension (73). Recent evidence suggests that plasma membrane domains such clathrin coated pits and caveolae as well as lipid metabolism are regulated by and provide feedback to YAP/TAZ (74, 75). Thus, additional insight will be gained by experiments and models that explore the role of membrane trafficking, membrane surface roughness, and membrane lipid metabolism on regulating YAP/TAZ mechanotransduction.

We note that our model, even though validated against data from the literature, is limited to certain components of the signaling pathway and that integration of other biochemical pathways will be important to build on this present work (76–80). For example, here we have only discussed the interaction between the ECM and single cells, yet cell-cell interactions are very important *in vivo* and may be responsible for some of the changes observed between 2D and 3D systems *in vitro (81, 82)*. Cell-cell and cell matrix adhesions can have synergistic and antagonistic effects (4), and regulation of YAP/TAZ by cell-cell communication through LATs and Merlin are additional key features to explore in future work. Incorporating cell type specific protein expression levels, realistic cellular geometries and physiologically relevant activation areas such as number, size, and localization of focal adhesions (83–88) will also offer further insight into how YAP/TAZ translocation is mediated by substrate stiffness and cell shape. Likewise, the timescale of YAP/TAZ nuclear localization could be important (89). We anticipate that studies such as this will empower mechanistic and systems exploration in cellular mechanotransduction, regenerative medicine, and tissue engineering.

## Author contributions

P.R. and S.F. conceived the project, analyzed the data, and wrote the manuscript. K.E.S constructed the initial versions of the model, the cellular and nuclear geometries and ran simulations for the well-mixed model and made Fig. 2. P.R. conducted all the final simulations and P.R and S.F. made all the figures of the manuscript. All authors agreed on the contents of the manuscript.

## Acknowledgments

We would like to thank Miriam Bell, Sural Ranamukhaarachchi, Allen Leung, and Will Leineweber for critical reading of the manuscript and their feedback.

## Funding

This work was supported by an NSF CAREER Award to S.I.F. (1651855) and an NSF GRFP Award to K.E.S. The Virtual Cell is supported by NIH Grant R24 GM137787 from the National Institute for General Medical Sciences.

## Supplementary Material

### A comment on length scale and conversions from well-mixed to spatial models

When converting between a well-mixed ordinary differential equation (ODE) model, such as those utilized in Sun et al. (1) and a partial differential equation (PDE) with boundary conditions, such as those used in the spatial model here, volume equations can become boundary fluxes. In order to convert the ODE to PDE, we need to use a length scale (n [micrometers]) to convert from volume units to the appropriate flux units. In our model, we use the volume to surface area ratio of the respective compartment, ex. cytoplasm or nucleus. However, to have the most realistic length scale possible, we utilize *n*_*r*_, which is a length scale taken from the corresponding compartment in a realistic geometry in a similar method to previous work (2). We define the length scale for converting between the volume of the cell to the volume in the nucleus from Table 2. By using a constant length scale 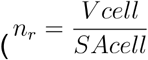, 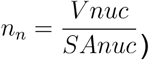, we are effectively defining a constant density of flux across all simulations. Based on how we defined various fluxes, we use this conversion for the cytoplasm and nucleus including the conversion of stiffness activation of FAK which occurs at the plasma membrane, and the flux of active and inactive forms of RhoA.

#### Sensitivity analysis method

The local sensitivities are defined as the change in the model output, Δ[*X*], caused by perturbation of a particular parameter, *α*, divided by the magnitude of the perturbation, *δα*. The local sensitivity is the derivative δ[*X*]/*δα* evaluated at a particular point in the parameter space. In this case, we utilize the normalized or scaled sensitivity given as,

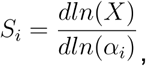

in order to compare the sensitivity across parameters. We derive the sensitivities of the steady-state fluxes of reactions with respect to the reaction rates in the system. 3.5 kPa is the center of the linear region of the slope in the YAP/TAZ Nuc/Cyto ratio in Fig. 2E. 0.1 and 5.7 kPa are close to the normal and tumorigenic stiffness respectively. The sensitivity values give information about the steady state of the parametric sensitivity coefficients for the system at these stiffnesses. The variable of interest is said to be robust with respect to a parameter if the log sensitivity is of order 1 (3). The variation (delta factor) used was 0.001 with a delta minimum of 1 × 10^−12^. This delta factor and minimum have been used previously (4). Within COPASI, we used the standard evolutionary programming method for 200 generations with a 20 population size using 1 random number generator to estimate all parameters. Goodness of fit between the experimental values and model output was determined using root-mean squared error (RMSE) and coefficient of determination, R^2^.

## Model geometry

The simulated geometry is one quarter of the cell as it is rotated around the z-axis. In order to define a realistic but theoretical cell shape in 3D, we utilized already existing geometry from Virtual Cell (5). The profile of the cell is defined using the following expression

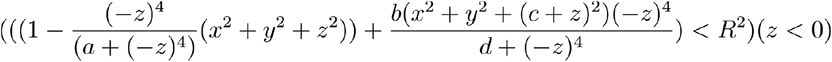

Here the radius of the cell at the widest point can be explicitly controlled by changing R. The height of the profile where x,y are small is primarily controlled by a combination of a, b and c. The height of the tail at the thinnest region where x,y are large can be controlled primarily by changing parameter d. See Table S8 for the sizes of the geometries used in each simulation. Volume is conserved across all simulation shapes. In these geometries we maintained a height greater than 0.2 *μm* in the lamellipodia according to (6). We model the nucleus as an ellipsoid for computational ease (7). In addition, we modeled the center of the nucleus in the center of the cell (8). We conducted simulations using four compartments: cytoplasm, plasma membrane (PM), nuclear membrane (NM), and nucleus (Table S7). We assume that all molecules in the model have sufficiently large molecular concentrations and model the reaction-diffusion equations using deterministic approaches.

We evaluate the steady state response to stiffness and scan over values of stiffness ranging from healthy physiologic range (0.1 kPa) (9), through tumor stiffness (5.7 kPa) (9), and up to glass substrates (70 GPa) (10).

### Boundary conditions for substrate dimensionality

In the spatial models, where the substrate stiffness was only presented at the bottom surface of the cell (2D and 2.xD), we used Virtual Cell’s built in Boolean operator to implement this condition. Essentially, we allowed for the substrate stiffness dependent FAK activation only on the bottom surface by multiplying that reaction with a Boolean operator for Z position. For the 3D stimulus, we allowed for the substrate stiffness dependent FAK activation on the whole surface. When testing the role of substrate presentation area versus total PM area in Figures 5, 6, for 2D stimulus, RhoA-GTP reactions were limited to the bottom surface using the position Boolean operator. For 2.xD and 3D conditions, RhoA-GTP reactions were allowed on the entire boundary surface of the cell.

## Supplementary Tables

**Table S1:** List of reactions and mass conservation relationships in the compartmental model. In this model, we used a series of simplified reactions in order to address the complex relationships between proteins, substrate stiffness, and cytosolic properties. We represent these relationships as reactions using both mass-action and Michaelis-Menten kinetics. In Table S1 we list the reaction and any modifier. We define a modifier as a protein which plays a role in the reaction as a catalyst but not as a reactant or a product. We set E to be a constant stiffness ranging from 0.1 to 70 × 10^6^ kPa and evaluate the steady state.

| List of Reactions | Modifier | Mass conservation |
| --- | --- | --- |
| **Module 1: Activation of FAK cascade due to stiffness** |  |  |
| $FAK \xrightleftharpoons[E]{E} pFAK$ | $E$ | $[FAK] + [pFAK] = [FAK_T]$ |
| $RhoA_{GDP} \rightleftharpoons RhoA_{GTP}$ | $pFAK$ | $[RhoA_{GDP}] + [RhoA_{GTP}] = [RhoA_T]$ |
| $ROCK \rightleftharpoons ROCK_A$ | $RhoA_{GTP}$ | $[ROCK] + [ROCK_A] = [ROCK_T]$ |
| $mDia \rightleftharpoons mDia_A$ | $RhoA_{GTP}$ | $[mDia] + [mDia_A] = [mDia_T]$ |
| **Module 2: Activation of downstream cascade** |  |  |
| $Myo \rightleftharpoons Myo_A$ | $ROCK_A$ | $[Myo] + [Myo_A] = [Myo_T]$ |
| $LIMK \rightleftharpoons LIMK_A$ | $ROCK_A$ | $[LIMK] + [LIMK_A] = [LIMK_T]$ |
| $Cofilin_P \xrightleftharpoons[LIMK_A]{LIMK_A} Cofilin_{NP}$ | $LIMK_A$ | $[Cofilin_P] + [Cofilin_{NP}] = [Cofilin_T]$ |
| $G_{actin} \rightleftharpoons F_{actin}$ | $mDia_A$<br>$Cofilin_{NP}$ | $[G_{actin}] + [F_{actin}] = [Actin_T]$ |
| **Module 3: Activation and translocation of YAP/TAZ** |  |  |
| $YAP_{TAZ_p} \rightleftharpoons YAP_{TAZ}$ | $F_{actin},$<br>$Myo_A$ | See below |
| $LaminA_p \xrightleftharpoons[E_{cytosol}]{E_{cytosol}} LaminA$ | $E_{cytosol}$ | $[LaminA_p] + [LaminA] = [LaminA_T]$ |
| $NPC \rightleftharpoons NPC_A$ | $LaminA,$<br>$F_{actin},$<br>$Myo_A$ | $[Cofilin_P] + [Cofilin_{NP}] = [Cofilin_T]$ |
| $YAP_{TAZ} \rightleftharpoons YAP_{TAZ_{nuc}}$ | $NPC_A$ | $([YAP_{TAZ_p}] + [YAP_{TAZ}]) * V_{cell} + [YAP_{TAZ_{nuc}}] * V_{nuc} = Constant$ |

**Table S2:** Reaction Rate and ODEs in Compartmental model.

| # | Rates | ODE | Rate ref |
| --- | --- | --- | --- |
| Module 1: Activation of FAK cascade due to stiffness |  |  |  |
| 1 | $R_1 = k_f * FAK + k_{sf} \frac{E}{C + E} [FAK]$ | $\frac{d[FAK]}{dt} = -R_2 - R_1$ | (1) |
| 2 | $R_2 = -k_{df} [pFAK]$ | $\frac{d[pFAK]}{dt} = R_2 + R_1$ | (1) |
| 3 | $R_3 = k_{fkp} (\gamma [pFAK]^n + 1) [RhoA_{GDP}]$ | $\frac{d[RhoA_{GDP}]}{dt} = -R_3 - n_r^{-1} * R_4$ | (1) |
| 4 | $R_4 = -k_{dp} [RhoA_{GTP}]$ | $\frac{d[RhoA_{GTP}]}{dt} = n_r * R_3 + R_4$ | (1) |
| 5 | $R_5 = -k_{drock} [ROCK_A]$ | $\frac{d[ROCK]}{dt} = -R_5 - n_r^{-1} * R_6$ | |
| 6 | $R_6 = k_{rp} [RhoA_{GTP}] [ROCK]$ | $\frac{d[ROCK_A]}{dt} = R_5 + n_r^{-1} * R_6$ | (1) |
| Module 2: Activation of downstream cascade |  |  |  |
| $T([ROCK_A]) = (\tanh(sc1 * ([ROCK_A] - ROCKB)) + 1) [ROCK_A] / 2$ | | | |
| 7 | $R_7 = -k_{dmDia} [mDia_A]$ | $\frac{d[mDia]}{dt} = -R_7 - n_r^{-1} * R_8$ | (1) |
| 8 | $R_8 = k_{mp} [RhoA_{GTP}] [mDia]$ | $\frac{dmDia_A}{dt} = R_7 + n_r^{-1} * R_8$ | (1) |
| 9 | $R_9 = k_{mr} (\epsilon T([ROCK_A]) + 1) [Myo] - k_{dmy} [Myo_A]$ | $\frac{d[Myo]}{dt} = -R_9$<br>$\frac{d[Myo_A]}{dt} = R_9$ | (1) |
| 10 | $R_{10} = k_{lr} (\tau T([ROCK_A]) + 1) [LIMK] - k_{dl} [LIMK_A]$ | $\frac{d[LIMK]}{dt} = -R_{10}$<br>$\frac{d[LIMK_A]}{dt} = R_{10}$ | (1) |
| 11 | $R_{11} = k_{turn-over} [Cofilin_P] - \frac{k_{catCofilin} [LIMK_A] [Cofilin_{NP}]}{k_m Cofilin + [Cofilin_{NP}]}$ | $\frac{d[Cofilin_P]}{dt} = -R_{11}$ | Forward<br>(1) |
| | | $\frac{d[Co\text{filin}_{NP}]}{dt} = R_{11}$ | Reverse (11) |
| 12 | $R_{12} = k_{ra}(aT([mDia_A]) + 1)[Gactin] - (k_{dep} + k_{fcl}[Co\text{filin}_{NP}])[Factin]$ | $\frac{d[Gactin]}{dt} = -R_{12}$<br>$\frac{d[Factin]}{dt} = R_{12}$ | (1) |
| Module 3: Activation and translocation of YAP/TAZ |  |  |  |
| 13 | $R_{13} = (k_{CN} + k_{CY}[Factin][Myo_A])[YAPT\text{AZ}_P] - k_{NC}[YAPT\text{AZ}]$ | $\frac{d[YAPT\text{AZ}_P]}{dt} = -R_{13}$<br>$\frac{d[YAPT\text{AZ}]}{dt} = R_{13} - n_n^{-1}R_{16}$ | (1) |
| 14 | $R_{14} = k_{fl}\frac{E_{cytosol}}{C_{laminA} + E_{cytosol}}[LaminA_p] - k_{rl}[LaminA]$<br>$E_{cytosol} = p[F_{cyto}]^{2.6}$<br>Cytosolic stiffness was modeled as a function of F-actin (see Notes) | $\frac{d[LaminA_p]}{dt} = -R_{14}$<br>$\frac{d[LaminA]}{dt} = R_{14}$ | This paper |
| 15 | $R_{15} = k_{fNPC}[LaminA][Factin][Myo_A][NPC] - k_r[NPC_A]$ | $\frac{d[NPC]}{dt} = -R_{15}$<br>$\frac{d[NPC_A]}{dt} = R_{15}$ | This paper |
| 16 | $R_{16} = (kin_b + k_{in} * [NPC_A])[YAPT\text{AZ}] - k_{out} * [YAPT\text{AZ}_{nuc}]$ | $\frac{d[YAPT\text{AZ}_{nuc}]}{dt} = n_n^{-1} * R_{16}$ | This paper |

Michaelis-Menten kinetics are used to model enzyme-catalyzed reactions. When a reaction is catalyzed by an enzyme with kinetic properties kcat and KM,

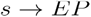

Then the product formation rate is given by

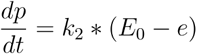

Under the T(ROCK approximation the rate of product formation is equivalent to the following:

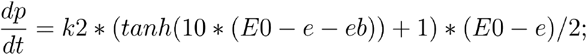

**Table S3:**
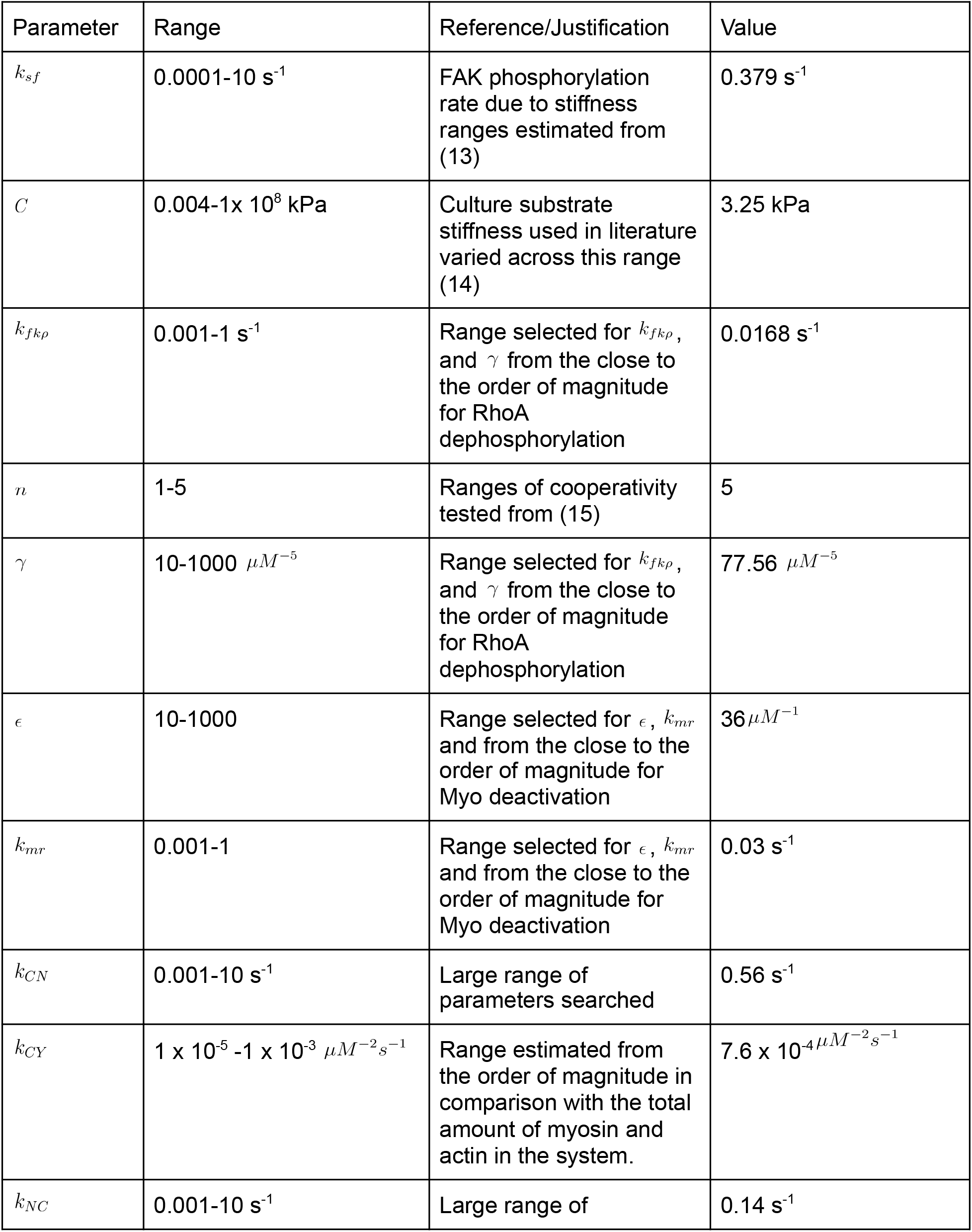
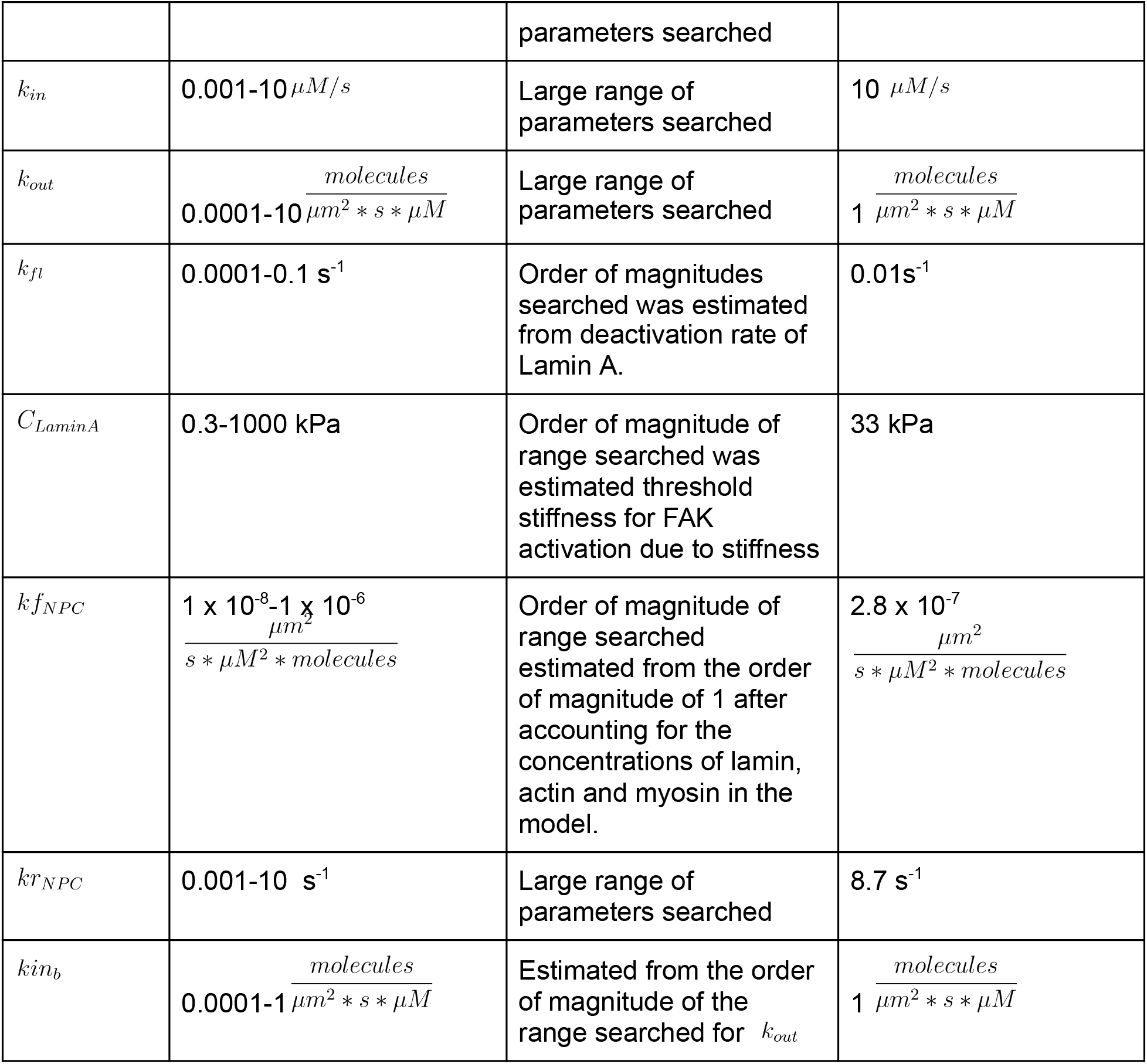
Ranges for parameter estimation. Of the parameters used in the model, 17 parameters were estimated using experimental data from the literature and using tools in COPASI to obtain best fits between models and experiments (12). In column 2 we report the range searched in COPASI for the parameter as well and in column 4 we report the final value found through the COPASI estimation process. Large ranges were searched for the value of each parameter.

**Table S4:**
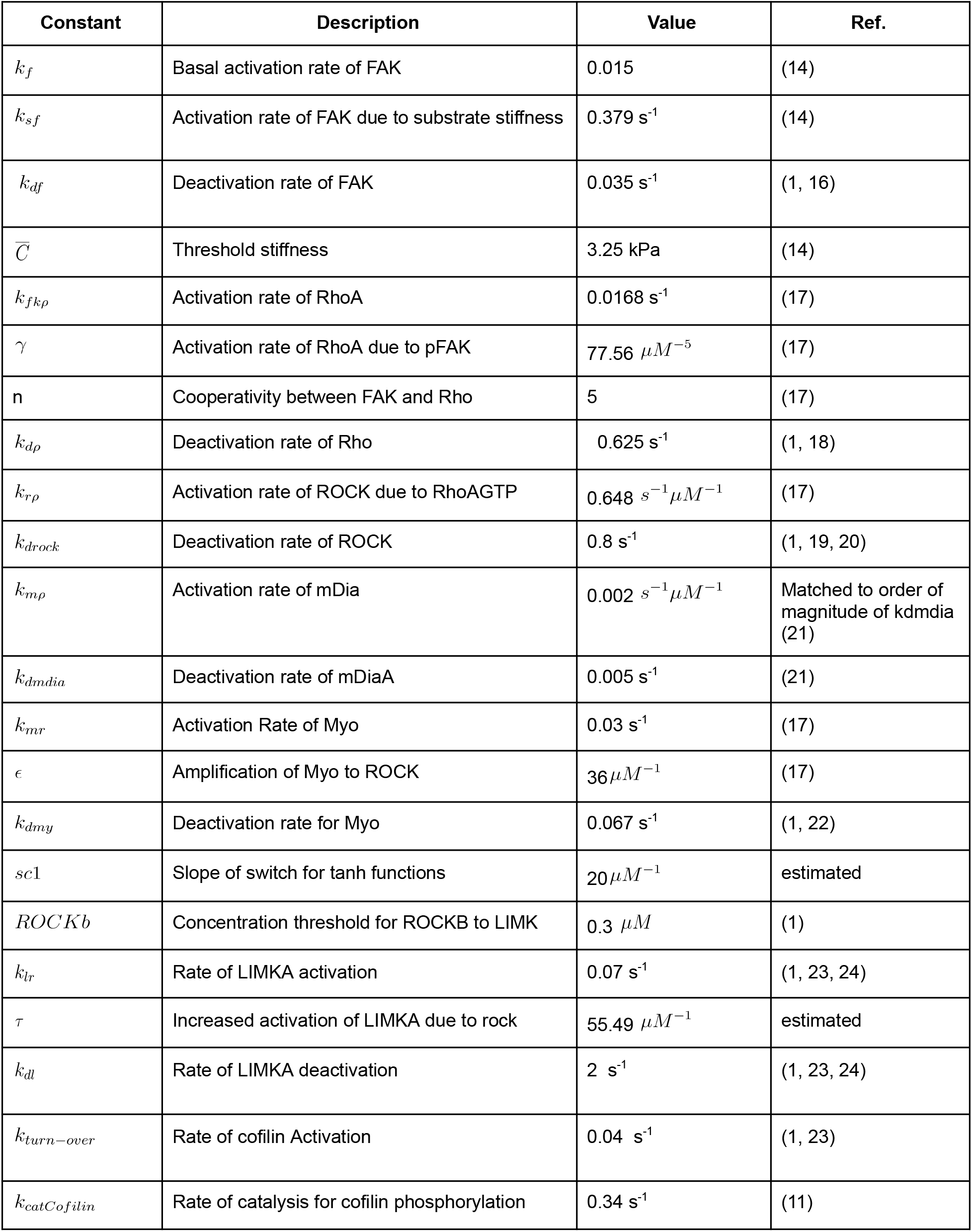
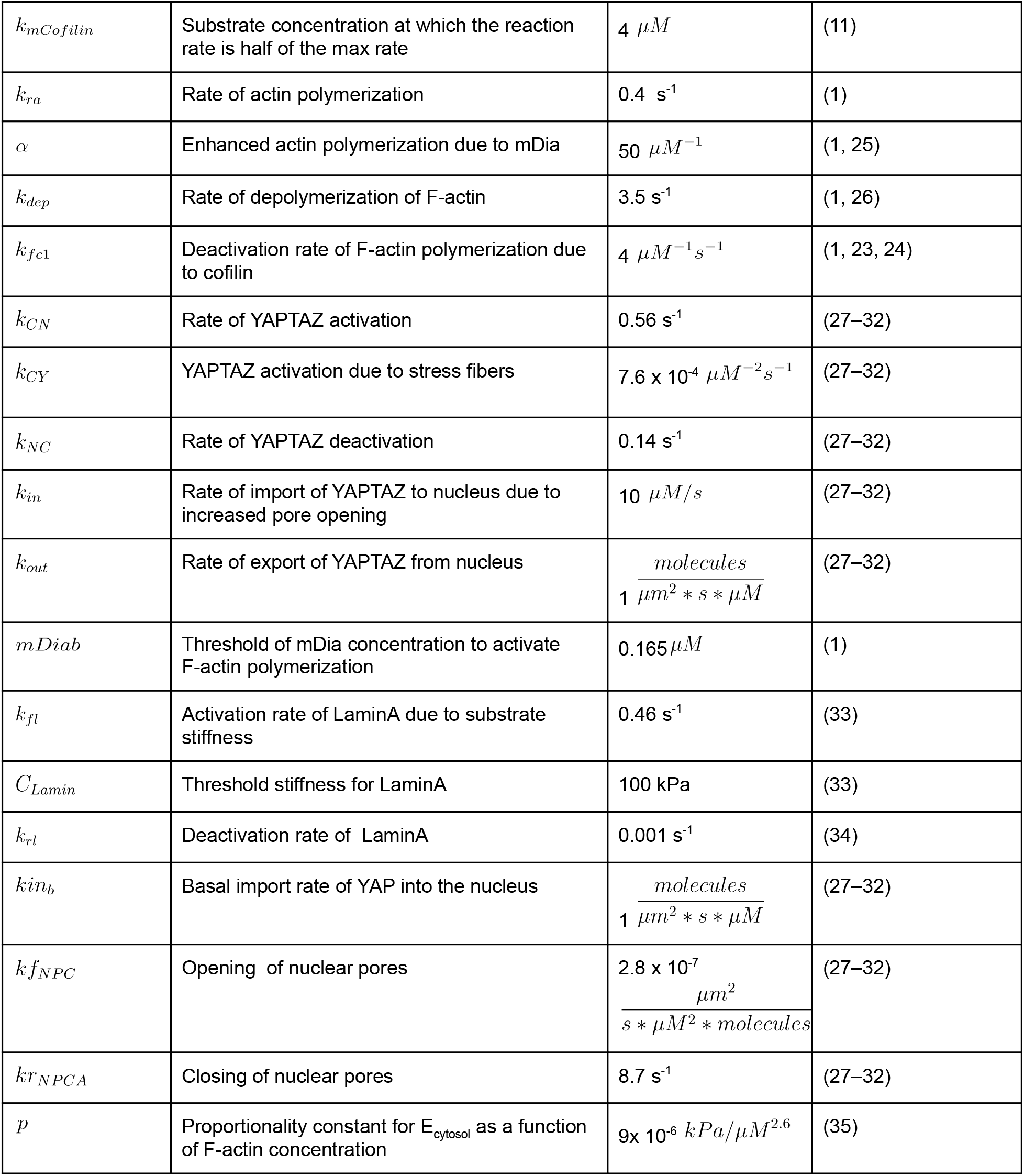
Parameters in compartmental model.

### Derivation of the relationship between F-actin and cytosolic stiffness

We derived the relationship between cytosolic stiffness and actin concentration from (35), as shown in Fig. S2A. Fig. S2B shows the model prediction over various substrate stiffnesses.The range of cytosolic stiffness is within those observed experimentally between (35, 36).

**Table S5:**
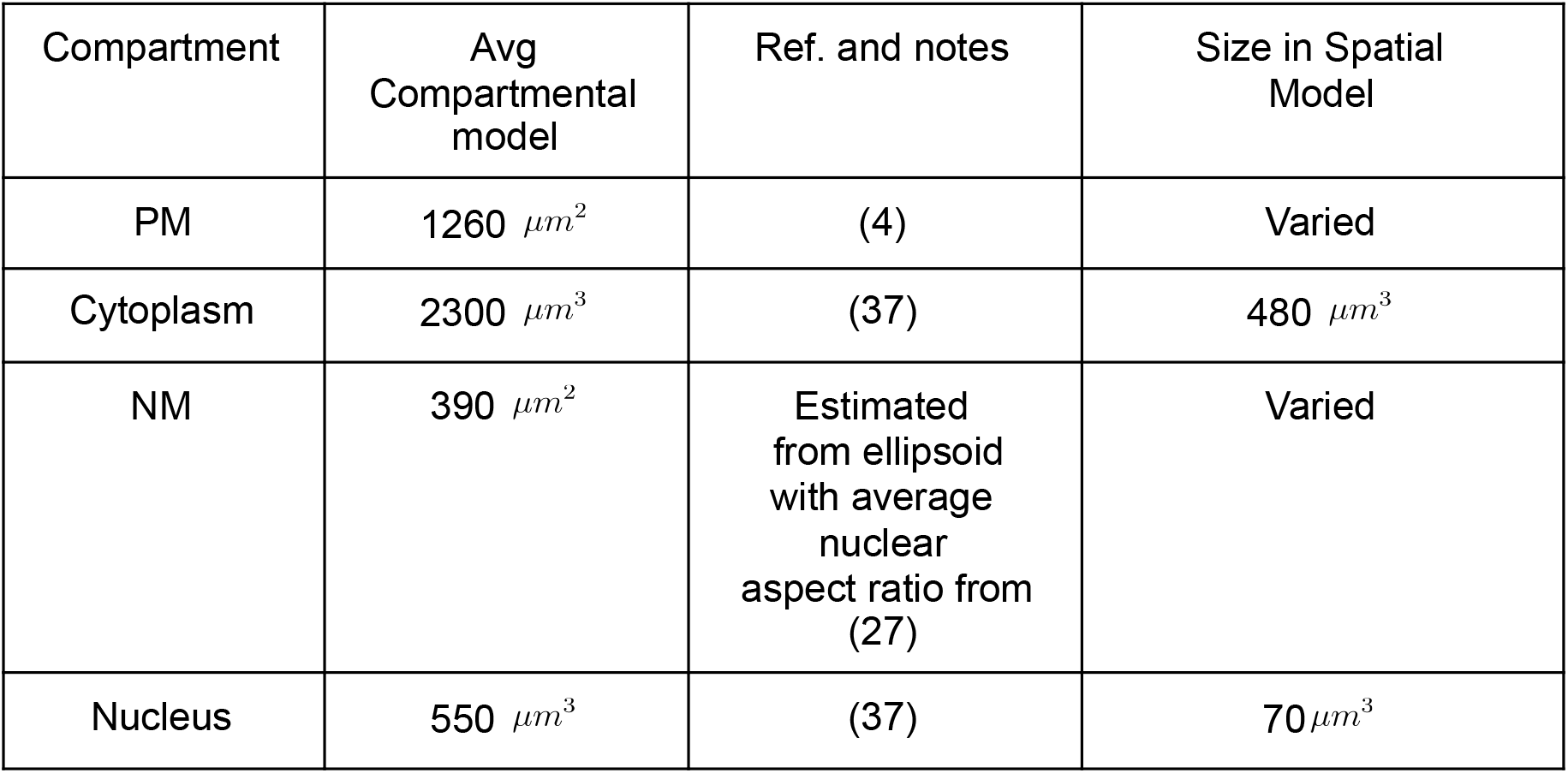
Sizes of different compartments used within the model.

| Compartment | Avg Compartmental model | Ref. and notes | Size in Spatial Model |
| --- | --- | --- | --- |
| PM | 1260 $\mu m^2$ | (4) | Varied |
| Cytoplasm | 2300 $\mu m^3$ | (37) | 480 $\mu m^3$ |
| NM | 390 $\mu m^2$ | Estimated from ellipsoid with average nuclear aspect ratio from (27) | Varied |
| Nucleus | 550 $\mu m^3$ | (37) | 70 $\mu m^3$ |

**Table S6:**
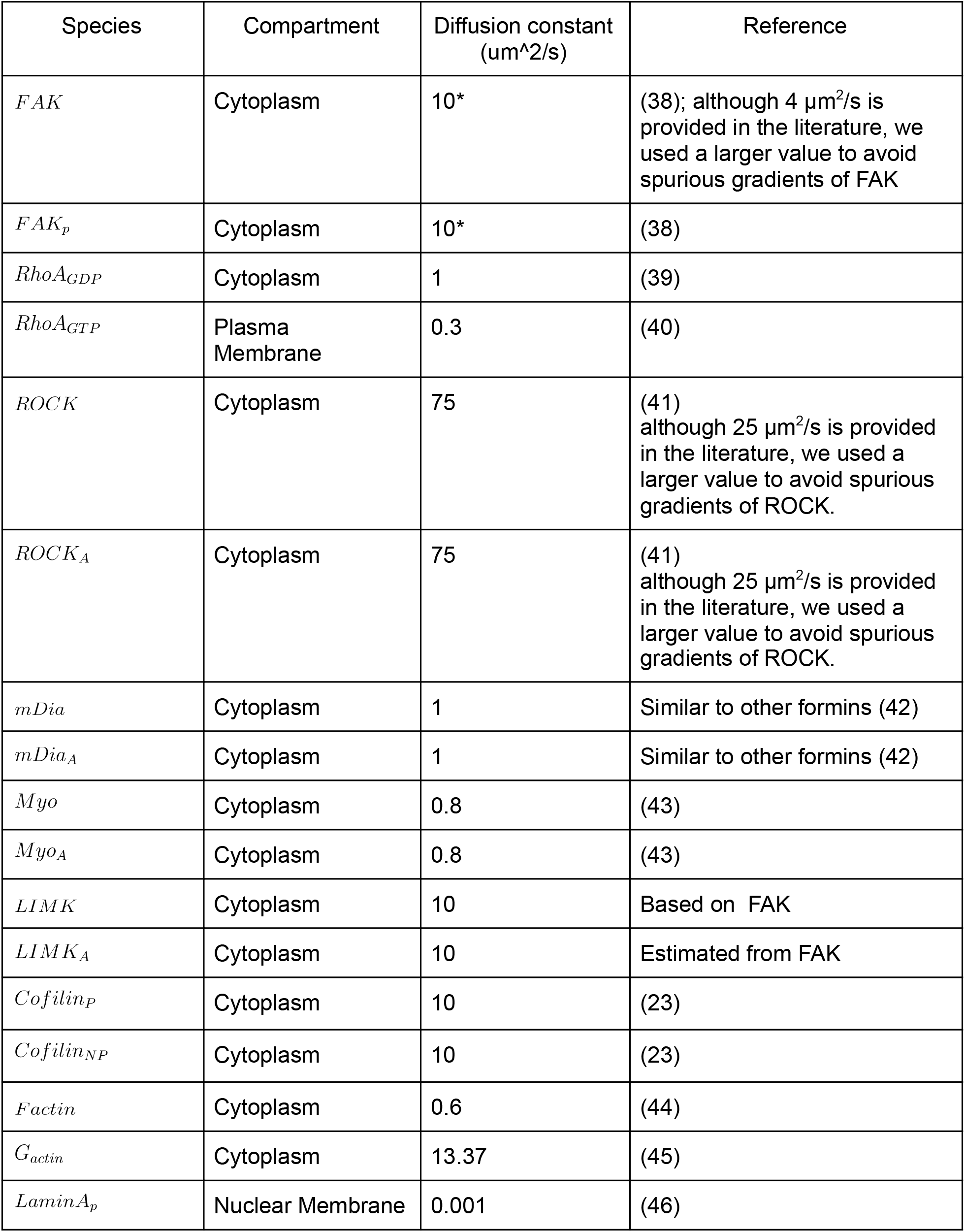
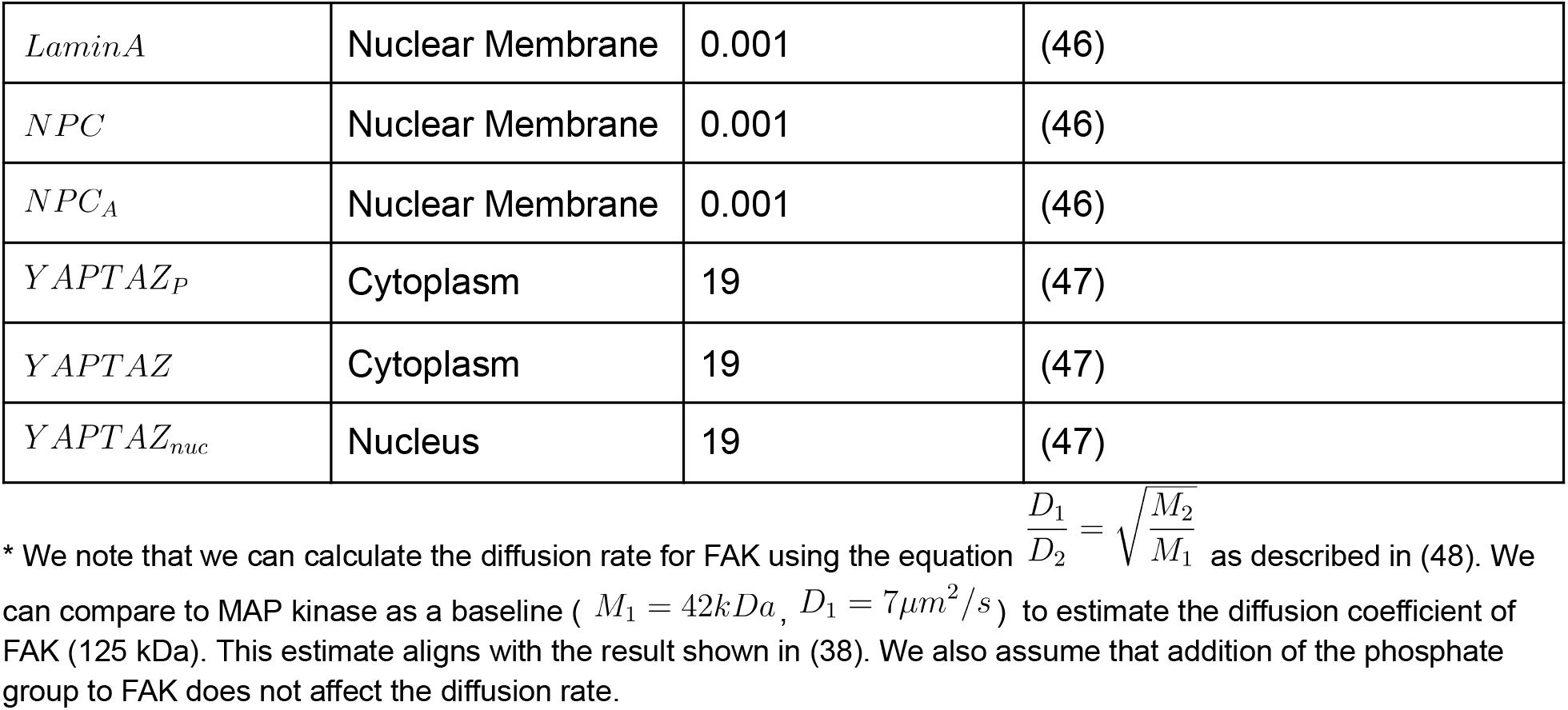
Diffusion Coefficients and Geometric Constraints.

**Table S7:** Total Concentrations of proteins per cell.

| Species | Amount | Reference |
| --- | --- | --- |
| *Fak<sub>T</sub>* | 1 $\mu\text{M}$ | Equivalent to concentration of RhoA as shown in (49) |
| *RhoA<sub>T</sub>* | 1 $\mu\text{M}$ | (11) |
| *ROCK<sub>T</sub>* | 1 $\mu\text{M}$ | (11) |
| *mDia<sub>T</sub>* | 0.8 $\mu\text{M}$ | Amount of mDia was approximately 80% of the total RhoA in a sample (50) |
| *Myo<sub>T</sub>* | 5 $\mu\text{M}$ | (11) |
| *LIMK<sub>T</sub>* | 2 $\mu\text{M}$ | (11) |
| *Cofilin<sub>T</sub>* | 2 $\mu\text{M}$ | (11) |
| *Actin<sub>T</sub>* | 500 $\mu\text{M}$ | (51) |
| *LaminA<sub>T</sub>* | 3500 *molecules/$\mu\text{m}^2$* | From (52), we estimate that the filament approximately consists of 5 molecules every 40 nm and ~3 protofilaments (3.5 nm diameter) for every intermediate filament (10 nm) (53). We measured the total length of the lamin filaments in a box, converted the total length to the total number of molecules (3*5/40 nm) and divided by the area of the box and repeated 4 times. This gives approximately 3500 +/- 600 molecules/$\mu\text{m}^2$ standard deviation. |
| *NPC<sub>T</sub>* | 6.5 *molecules/$\mu\text{m}^2$* | From (54) we calculate the total number from Fig 1 G, H, I (~2500) divided by the nuclear surface area used in the model geometry section<br>Similar to density found in (55) |
| *YAPTAZ<sub>T</sub>* | 1 $\mu\text{M}$ | Similar levels YAP/TAZ and FAK (56) |

**Table S8:** Initial Concentrations for model.

| Species | Concentration after running 0 stimulus (E=0 kPa) |
| --- | --- |
| $Cofilin_{NP}$ | $1.8 \mu M$ |
| $Cofilin_P$ | $0.2 \mu M$ |
| $FAK$ | $0.7 \mu M$ |
| $FAK_P$ | $0.3 \mu M$ |
| $F_{actin}$ | $17.9 \mu M$ |
| $G_{actin}$ | $482.4 \mu M$ |
| $LaminA$ | $0.0 \text{ molecules}/\mu m^2$ |
| $LaminA_P$ | $3500.0 \text{ molecules}/\mu m^2$ |
| $LIMK$ | $1.9 \mu M$ |
| $LIMK_A$ | $0.1 \mu M$ |
| $mDia$ | $0.8 \mu M$ |
| $mDia_A$ | $0.0 \mu M$ |
| $Myo$ | $3.5 \mu M$ |
| $Myo_A$ | $1.5 \mu M$ |
| $NPC$ | $6.5 \text{ molecules}/\mu m^2$ |
| $NPC_A$ | $0.0 \text{ molecules}/\mu m^2$ |
| $RhoA_{GDP}$ | $1.0 \mu M$ |
| $RhoA_{GTP}$ | $33.6 \text{ molecules}/\mu m^2$ |
| $ROCK$ | $1.0 \mu M$ |
| $ROCK_A$ | $0.0 \mu M$ |
| $YAPTAZ$ | $0.7 \mu M$ |
| $Y_{APT AZ_{nuc}}$ | $0.7 \mu M$ |
| $Y_{APT AZ_P}$ | $0.2 \mu M$ |

**Table S9:**
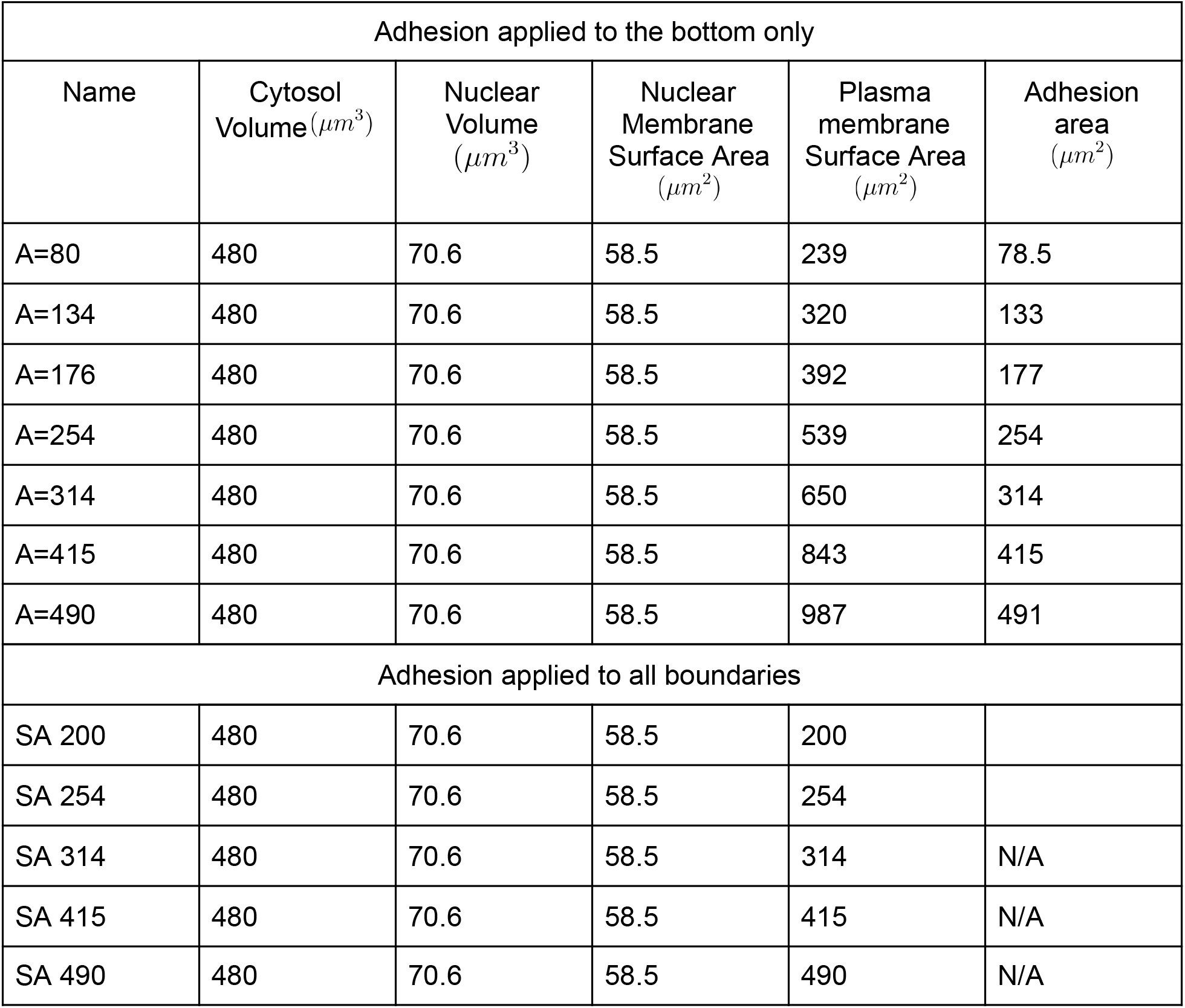
Parameters in the spatial model.

| Adhesion applied to the bottom only |  |  |  |  |  |
| --- | --- | --- | --- | --- | --- |
| Name | Cytosol Volume ($\mu m^3$) | Nuclear Volume ($\mu m^3$) | Nuclear Membrane Surface Area ($\mu m^2$) | Plasma membrane Surface Area ($\mu m^2$) | Adhesion area ($\mu m^2$) |
| A=80 | 480 | 70.6 | 58.5 | 239 | 78.5 |
| A=134 | 480 | 70.6 | 58.5 | 320 | 133 |
| A=176 | 480 | 70.6 | 58.5 | 392 | 177 |
| A=254 | 480 | 70.6 | 58.5 | 539 | 254 |
| A=314 | 480 | 70.6 | 58.5 | 650 | 314 |
| A=415 | 480 | 70.6 | 58.5 | 843 | 415 |
| A=490 | 480 | 70.6 | 58.5 | 987 | 491 |
| Adhesion applied to all boundaries |  |  |  |  |  |
| SA 200 | 480 | 70.6 | 58.5 | 200 |  |
| SA 254 | 480 | 70.6 | 58.5 | 254 |  |
| SA 314 | 480 | 70.6 | 58.5 | 314 | N/A |
| SA 415 | 480 | 70.6 | 58.5 | 415 | N/A |
| SA 490 | 480 | 70.6 | 58.5 | 490 | N/A |

**Table S10:** Vcell simulations for each figure.

|  |  |
| --- | --- |
| Figure 3: Spatial simulations comparing substrate stiffness presentation<br>Published Model Name: to be filled out after revisions and acceptance of manuscript. |  |
| Application name | Simulation 1 |
| R15 2D Fig 3 and 4<br>Bottom -- 2D stimulus<br>presentation | R 15 Bot 0.1,5.7,70E6<br>simulation for 3 values<br>of E for geometry with radius 15 with stimulus from the bottom |
| R15 3D Figure 3-4 --<br>3D stimulus<br>presentation | R 15 All 0.1,5.7,70E6<br>simulation for 3 values<br>of E for geometry with radius 15 with stimulus from the bottom |
| Figure 4: Spatial simulations comparing substrate stiffness presentation for three different cell shapes<br>Published Model Name: to be filled out after revisions and acceptance of manuscript. |  |
| R10 2D Figure 4<br>Bottom -- 2D stimulus<br>presentation | R 10 Bot 0.1,5.7,70E6<br>simulation for 3 values<br>of E for geometry with radius 10 with stimulus from the bottom |
| R10 3D Figure 4 -- 3D<br>stimulus presentation | R 10 All 0.1,5.7,70E6<br>simulation for 3 values<br>of E for geometry with radius 10 with stimulus from the bottom |
| R13 2D Figure 4<br>Bottom -- 2D stimulus<br>presentation | R 13 Bot 0.1,5.7,70E6<br>simulation for 3 values<br>of E for geometry with radius 13 with stimulus from the bottom |
| R13 3D Figure 4 -- 3D<br>stimulus presentation | R 13 All 0.1,5.7,70E6<br>simulation for 3 values<br>of E for geometry with radius 13 with stimulus from the bottom |
| R15 2D Fig 3 and 4<br>Bottom -- 2D stimulus<br>presentation | R 15 Bot 0.1,5.7,70E6<br>simulation for 3 values<br>of E for geometry with radius 15 with stimulus from the bottom |
| R15 3D Figure 3-4 --<br>3D stimulus<br>presentation | R 15 All 0.1,5.7,70E6<br>simulation for 3 values<br>of E for geometry with radius 15 with stimulus from the bottom |
| Figures 5 and 6: Spatial simulations comparing substrate stiffness presentation for three different cell shapes<br>Published Model Name: to be filled out after revisions and acceptance of manuscript.<br>Two different model files were developed. 2D and 3D simulations can be found in Filename XXX while 2.XD<br>simulations can be found in YYY. |  |
| SA 200 2.xD Figure 5,6 | R 16 All 0.1,5.7,70E6 |
|  | Simulation for 3 values of E for geometry with radius 16 with stimulus from all boundaries |
| SA 200 3D Figure 5,6 | R 16 All 0.1,5.7,70E6<br>Simulation for 3 values of E for geometry with radius 16 with stimulus from all boundaries |
| SA 250 2.xD Figure 5,6 | R 18 All 0.1,5.7,70E6<br>Simulation for 3 values of E for geometry with radius 18 with stimulus from all boundaries |
| SA 250 3D Figure 5,6 | R 18 All 0.1,5.7,70E6<br>Simulation for 3 values of E for geometry with radius 18 with stimulus from all boundaries |
| SA 314 2.xD Figure 5,6 | R 20 All 0.1,5.7,70E6<br>Simulation for 3 values of E for geometry with radius 20 with stimulus from all boundaries |
| SA 314 3D Figure 5,6 | R 20 All 0.1,5.7,70E6<br>Simulation for 3 values of E for geometry with radius 20 with stimulus from all boundaries |
| SA 415 2.xD Figure 5,6 | R 23 All 0.1,5.7,70E6<br>Simulation for 3 values of E for geometry with radius 23 with stimulus from all boundaries |
| SA 415 3D Figure 5,6 | R 23 All 0.1,5.7,70E6<br>Simulation for 3 values of E for geometry with radius 23 with stimulus from all boundaries |
| SA 490 2.xD Figure 5,6 | R 26 All 0.1,5.7,70E6<br>Simulation for 3 values of E for geometry with radius 26 with stimulus from all boundaries |
| SA 490 3D Figure 5,6 | R 26 All 0.1,5.7,70E6<br>Simulation for 3 values of E for geometry with radius 26 with stimulus from all boundaries |
| 2D simulations are in a different model file named: YYY |  |
| SA 200 2D Figure 5,6 | R 16 All 0.1,5.7,70E6<br>Simulation for 3 values of E for geometry with radius 16 with stimulus from all boundaries |
| SA 250 2D Figure 5,6 | R 18 All 0.1,5.7,70E6<br>Simulation for 3 values of E for geometry with radius 18 with stimulus from all boundaries |
| SA 314 2 D Figure 5,6 | R 20 All 0.1,5.7,70E6<br>Simulation for 3 values of E for geometry with radius 20 with stimulus from all boundaries |
| SA 415 2D Figure 5,6 | R 23 All 0.1,5.7,70E6<br>Simulation for 3 values of E for geometry with radius 23 with stimulus from all boundaries |
| SA 490 2D Figure 5,6 | R 26 All 0.1,5.7,70E6<br>Simulation for 3 values of E for geometry with radius 26 with stimulus from all boundaries |
| Figure 7: Spatial simulations comparing substrate stiffness presentation for three different cell shapes and three different nuclear shapes<br>Published Model Name: to be filled out after revisions and acceptance of manuscript. |  |
| R10 LD 0.5 Figure 7 | R10 Bottom Tall 0.1,5.7,70E6<br>Nucleus with 0.5 times the L/D ratio to the control and the shape with radius 10<br>Here $k_{in}$ is multiplied by 0.5 as the shape factor. |
| R10 LD 1.5 Figure 7 | R10 Bottom Tall 0.1,5.7,70E6<br>Nucleus with 1.5 times the L/D ratio to the control and the shape with radius 10<br>Here $k_{in}$ is multiplied by 1.5 as the shape factor. |
| R13 LD 0.5 Figure 7 | R13 Bottom Tall 0.1,5.7,70E6<br>Nucleus with 0.5 times the L/D ratio to the control and the shape with radius 10<br>Here $k_{in}$ is multiplied by 0.5 as the shape factor. |
| R13 LD 1.5 Figure 7 | R13 Bottom Tall 0.1,5.7,70E6<br>Nucleus with 1.5 times the L/D ratio to the control and the shape with radius 13<br>Here $k_{in}$ is multiplied by 1.5 as the shape factor. |
| R15 LD 0.5 Figure 7 | R15 Bottom Tall 0.1,5.7,70E6 |
|  | Nucleus with 0.5 times the L/D ratio to the control and the shape with radius 15<br>Here $k_{in}$ is multiplied by 0.5 as the shape factor. |
| R15 LD 1.5 Figure 7 | R15 Bottom Tall 0.1,5.7,70E6<br>Nucleus with 1.5 times the L/D ratio to the control and the shape with radius 15<br>Here $k_{in}$ is multiplied by 1.5 as the shape factor. |
| R16 LD 0.5 Figure 7 | R10 Bottom Tall 0.1,5.7,70E6<br>Nucleus with 0.5 times the L/D ratio to the control and the shape with radius 10<br>Here $k_{in}$ is multiplied by 0.5 as the shape factor. |
| R16 LD 1.5 Figure 7 | R10 Bottom Tall 0.1,5.7,70E6<br>Nucleus with 1.5 times the L/D ratio to the control and the shape with radius 16<br>Here $k_{in}$ is multiplied by 1.5 as the shape factor. |

## Supplementary figures

**Figure S1.**
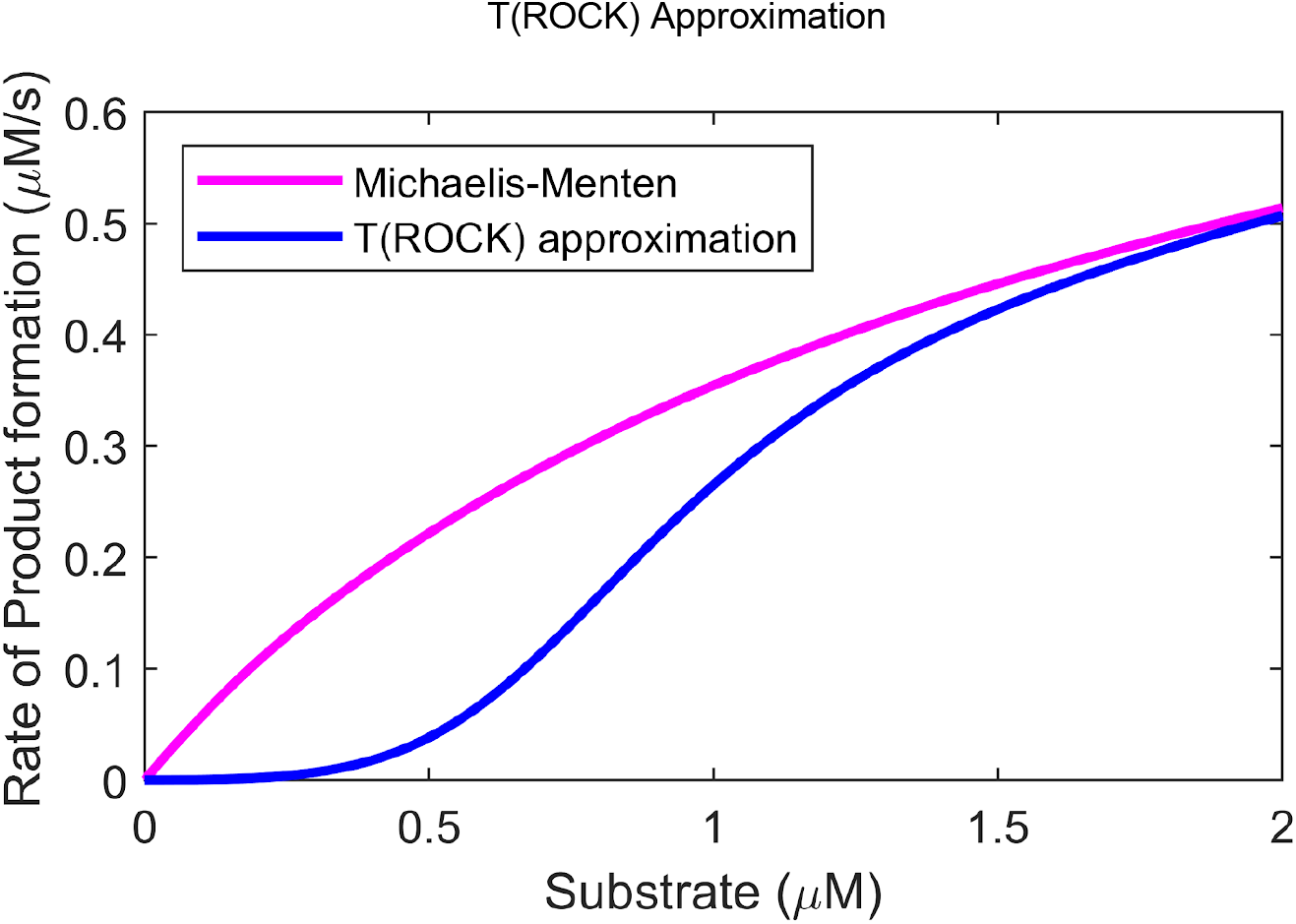
The T(ROCK) approximation approximates the Michaelis Menten curve but shows significantly more threshold effects for very low substrate concentrations.

**Figure S2.**
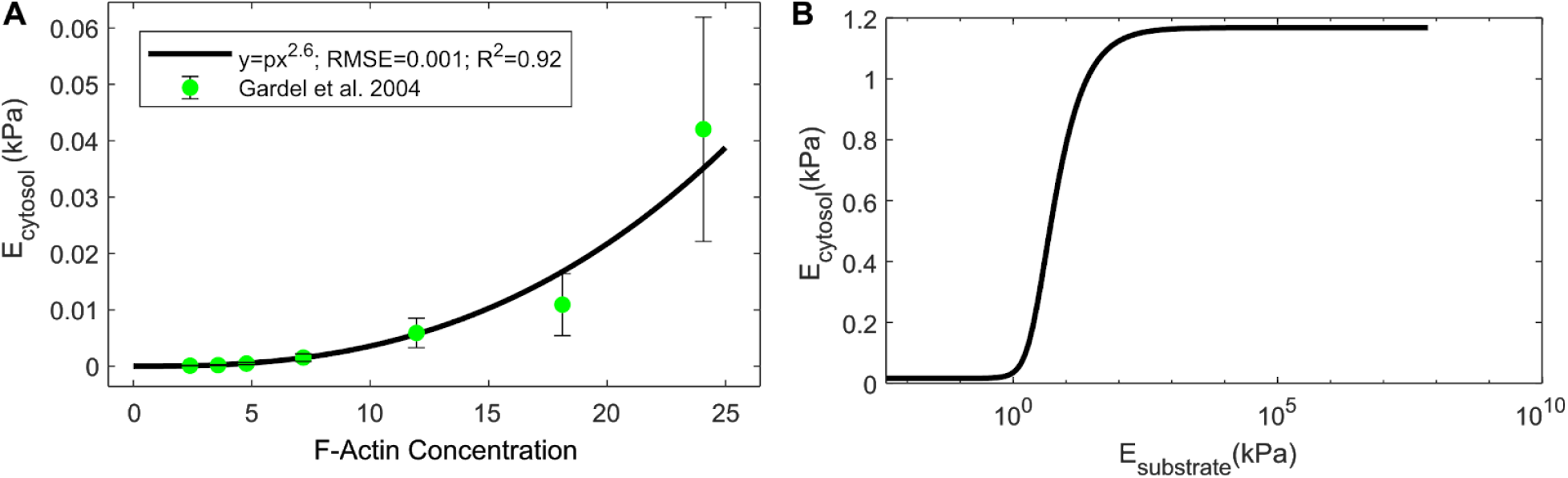
A) Relationship between F-actin concentration and substrate stiffness was fit from (35). B) Model prediction for cytosolic stiffness over substrate stiffness.

Sensitivity to full model parameters and initial conditions

**Figure S3.**
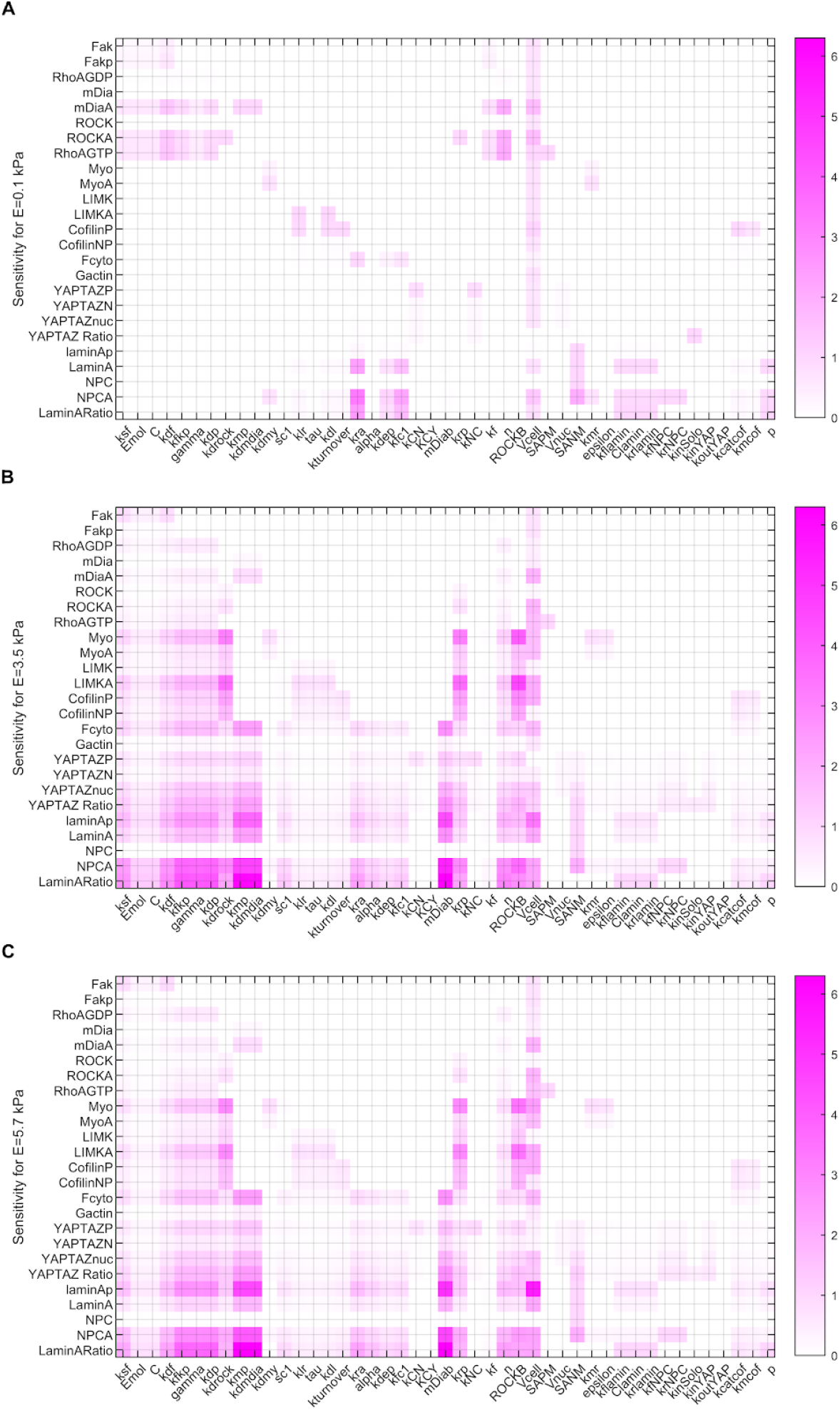
Sensitivity analysis for all of the variables and all of the parameters in the full model for 3 stiffnesses A) 0.1 kPa, B) 3.5 kPa and C) 5.7kPa.

**Figure S4.**
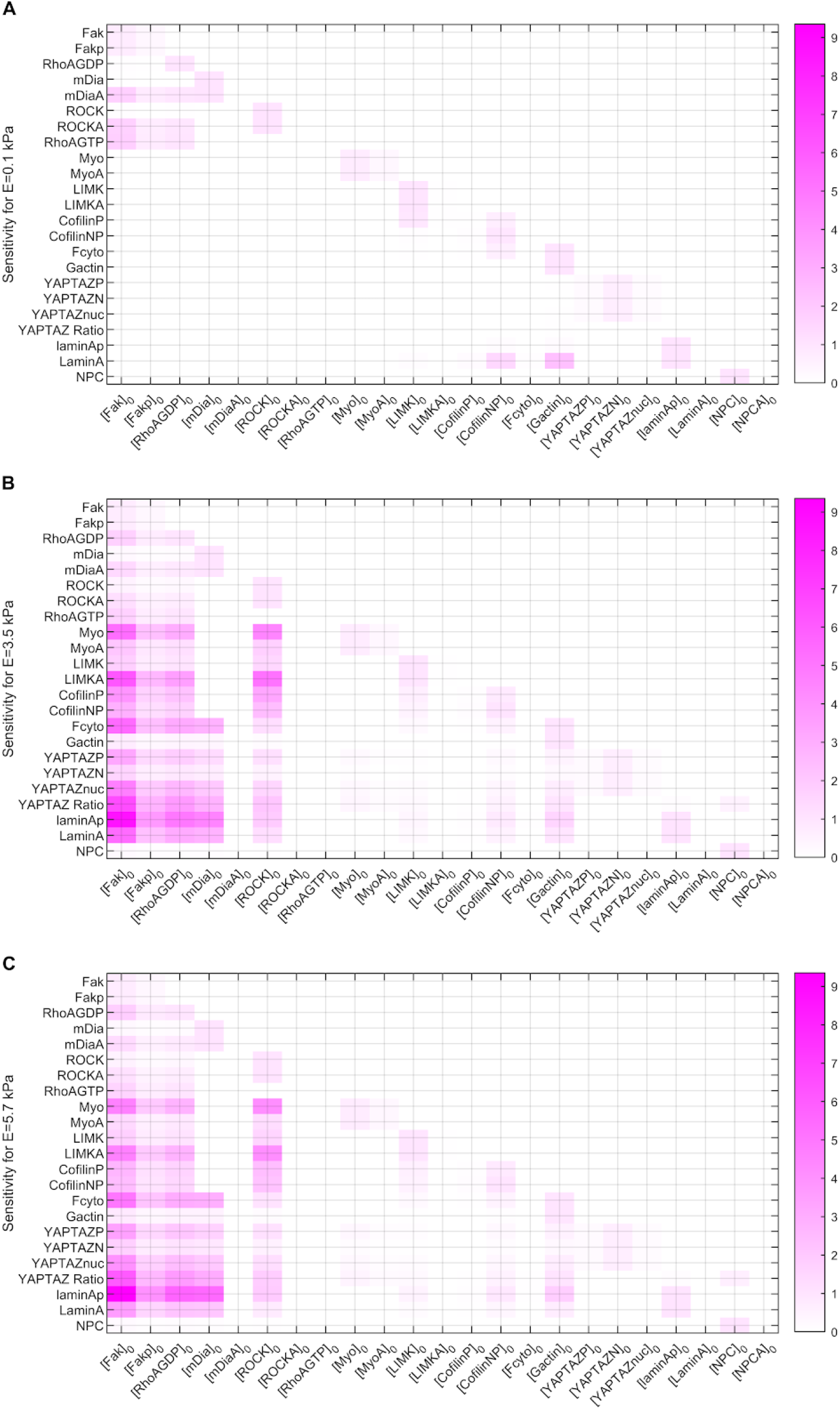
Sensitivity analysis for all of the initial conditions in the model A) 0.1 kPa, B) 3.5 kPa and C) 5.7 kPa.

**Figure S5.**
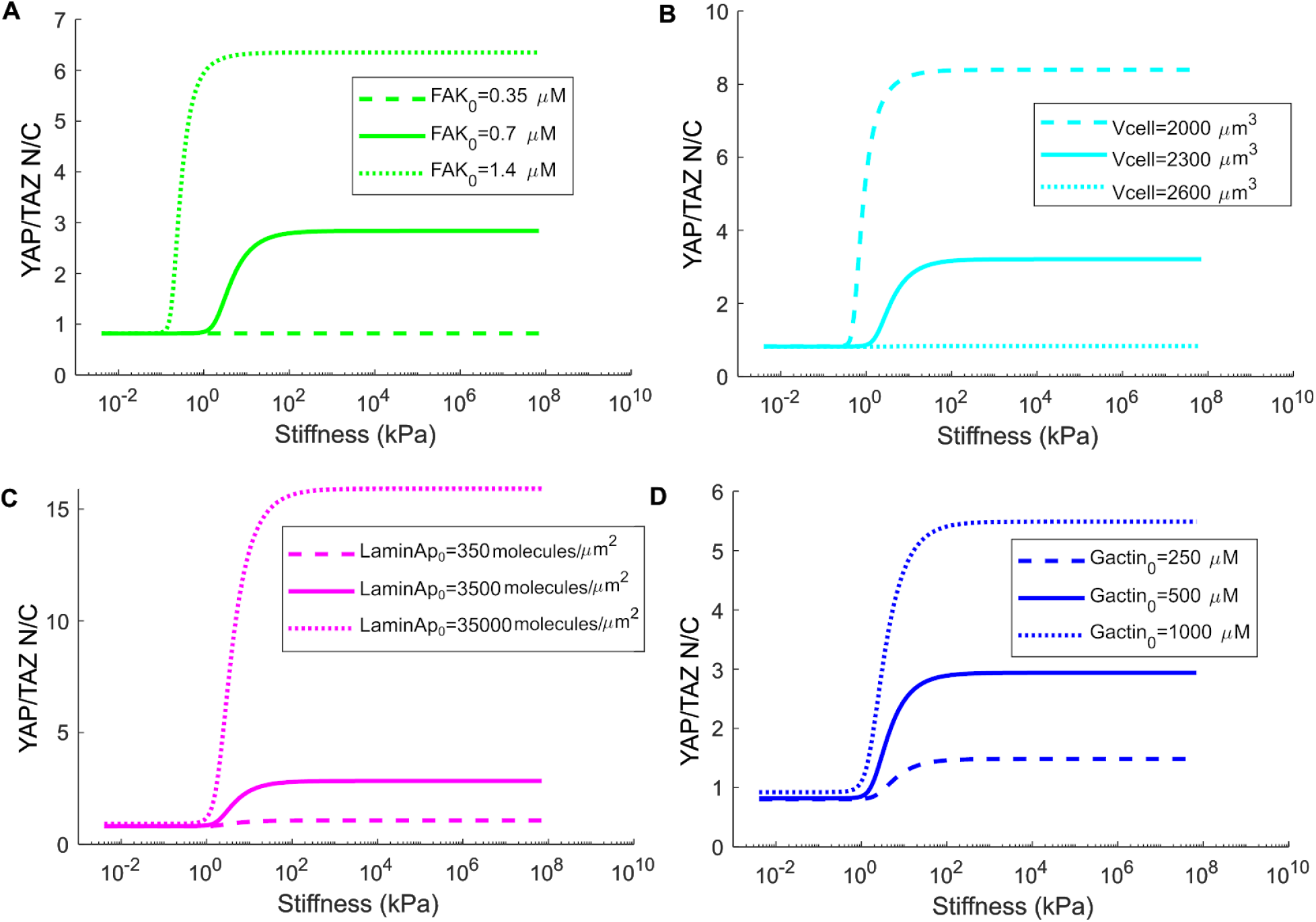
Model predictions for changes to selected parameters. (*A*) Effect of varying initial condition of FAK on the YAP/TAZ Nuc/Cyto Ratio. Initial condition used in the model is 0.7 *μM*. (*B*) Effect of varying volume of the cell in range from (37). (*C*) Model predictions for varying the initial conditions of Lamin A over a large range. The initial condition used in the model is 3500 *molecules/μm*^2^, which we found as described in Table S6. (*D*) Effect of varying actin initial condition across 2 orders of magnitude within physiologic concentration ranges from 250 *μM* (57) to ~1000 *μM* (58). The initial condition used in the model is 500 *μM* (51).

**Figure S6:**
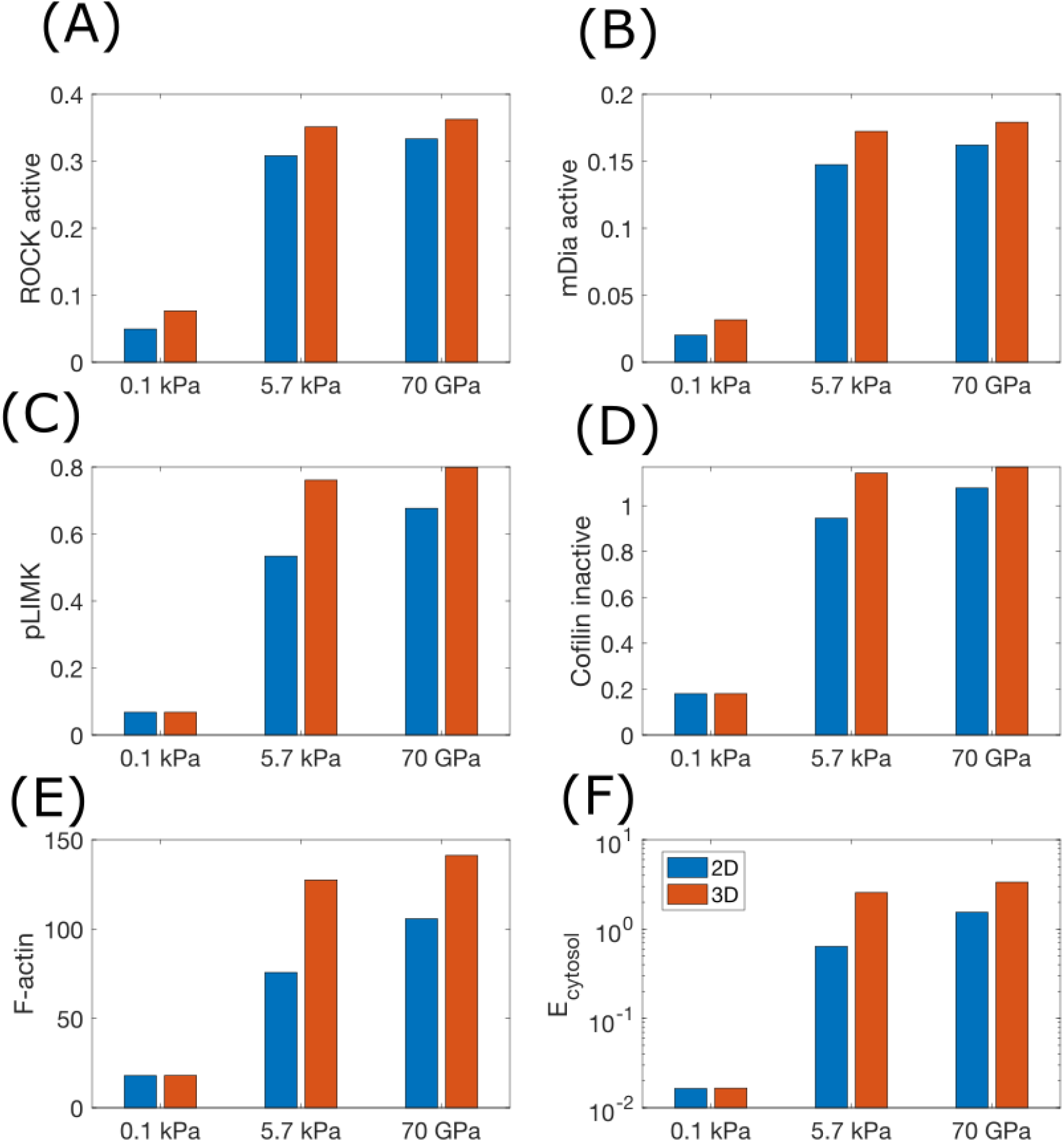
Comparison of 2D versus 3D substrate presentation for different stiffnesses. The geometry used is the same as Figure 3. (A) ROCK, (B) mDia, (C) LIMK active, (D) inactive cofilin (phospho-cofilin), (E) F-actin all in μM. (F) Cytosolic stiffness in kPa for these conditions. Note that the vertical axis of panel F is in the log scale.

**Figure S7:**
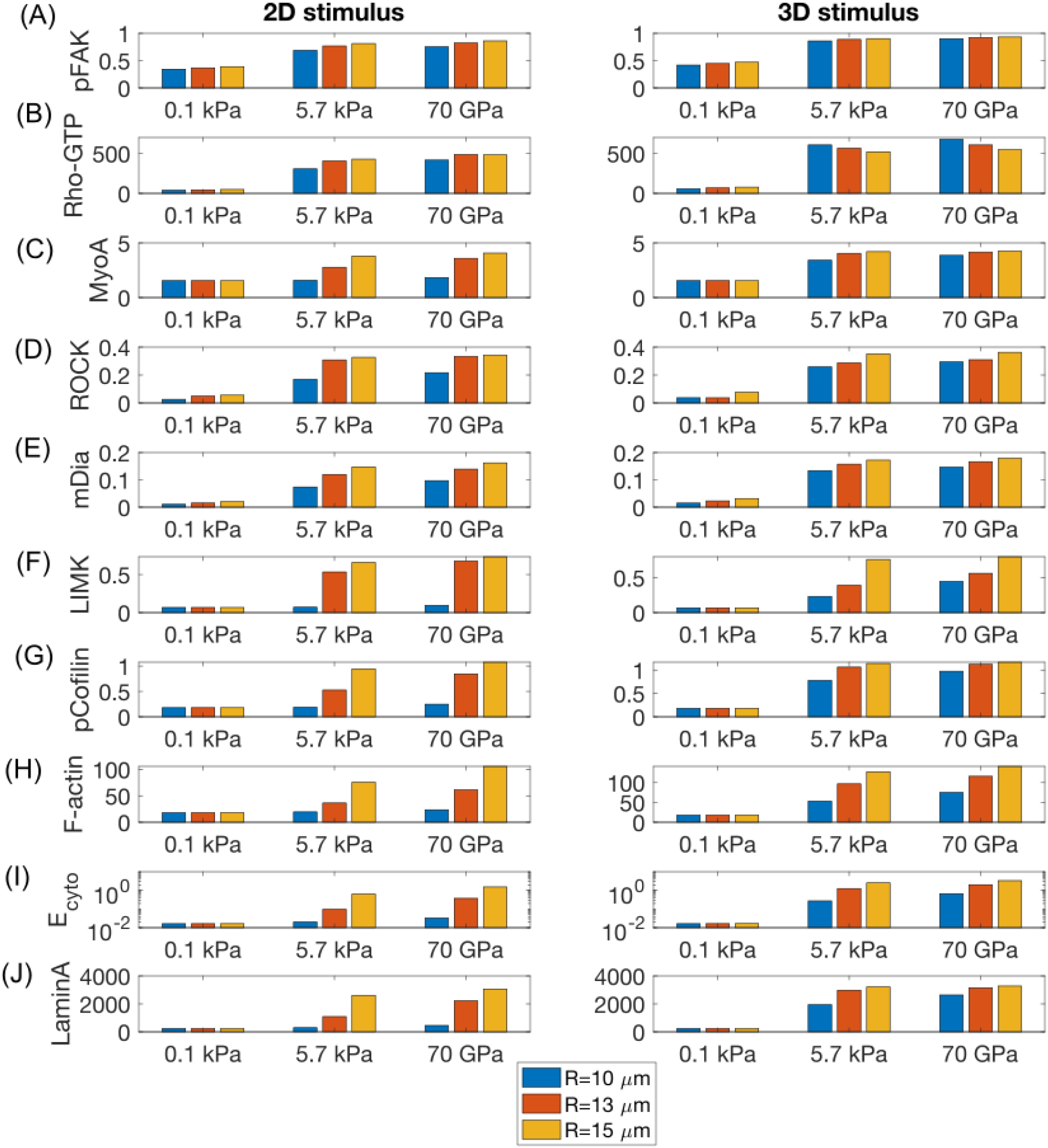
Steady state values (t=4000 s) of the different upstream species in Modules 1, 2, and 3 for different cell elongations and different stiffness. With the exception of cytosolic stiffness, Ecyto, all other vertical scales are linear scales.

## Notes

### Competing Interest Statement

The authors have declared no competing interest.

